# Integrating Acoustic, Prosodic, and Phonological Features for Automatic Alzheimer’s Detection

**DOI:** 10.64898/2026.01.16.699892

**Authors:** M. Zakaria Kurdi

## Abstract

Early and accurate diagnosis of Alzheimer’s Disease (AD) is critical for effective intervention. While previous studies have explored speech-based biomarkers for AD, this paper presents the first systematic investigation of acoustic, prosodic, and phonological speech features for detecting this neurocognitive disorder. Our study has two main objectives: (1) to assess the individual impact of AD on a wide range of speech features, and (2) to identify the most informative feature subsets using a combination of four feature ranking methods and seven machine learning classifiers.

We conducted our experiments using the publicly available ADReSS Challenge dataset, allowing for direct comparison with prior speech-based approaches. Our analysis focused on continuous acoustic and prosodic measures as well as discrete phonological features, both independently and in combination. The best performance, with an F1-score of 0.89, was achieved using the optimal subset of acoustic features with ensemble learning and by integrating all three feature types, suggesting that combining continuous and categorical speech features offers complementary diagnostic value.

This approach surpasses all previous speech-only methods in the ADReSS Challenge and comes close to the best reported overall result from the campaign’s dataset, even without using lexical information. The results were further validated through positive outcomes on the Delaware corpus (focused on MCI) and a set of speeches from President Ronald Reagan (who was diagnosed with Alzheimer’s). These findings suggest that speech patterns, beyond just the content, are more indicative of Alzheimer’s disease than previously thought, underscoring the potential of multi-layered speech analysis for non-invasive AD detection.

## 1. Introduction

Aging is accompanied by gradual cognitive, motor, and neurological changes that influence speech production. Older adults often exhibit reduced fluency, slower articulation, decreased articulatory precision, and prosodic flattening, patterns that reflect underlying declines in executive control, motor coordination, and neural efficiency (Linville, 2001). Because these alterations emerge naturally in spontaneous communication, speech offers a rich, unobtrusive window into neurocognitive health.

Alzheimer’s Disease (AD) is a major public health concern that disproportionately affects aging populations. According to the Alzheimer’s Association (2025), more than 7.2 million Americans live with AD. Early detection is critical for enabling timely intervention, which can delay cognitive decline and improve quality of life (Rasmussen & Langerman, 2019). However, existing diagnostic practices, such as neuroimaging, cerebrospinal fluid analysis, and lengthy cognitive assessments, are often invasive, costly, and difficult to scale, creating barriers to early screening and monitoring.

Speech has emerged as a promising, non-invasive digital biomarker for detecting cognitive impairment associated with AD. Subtle alterations in phonology, prosody, and acoustic quality may precede more evident cognitive symptoms, providing an early indicator of disease onset (Luchesi Cera et al., 2023; Koo et al, 2020; Haider et al, 2019; Szatloczki et al., 2015). Advances in machine learning (ML) have further enabled automated analysis of speech patterns, offering scalable and remote assessment solutions for clinical and telehealth settings (Fernández Montenegro et al, 2020). Nevertheless, many prior approaches have focused on limited features sets that has not been selected based on objective criteria, reducing diagnostic power and generalizability across languages and recording conditions.

In this study, we develop and validate an explainable machine learning framework that integrates a broad spectrum of speech features, including segmental (phonological), acoustic, and prosodic cues, to detect AD. This multidimensional design captures complementary aspects of speech production and is robust to linguistic and recording variability, making it particularly suitable for aging populations with diverse backgrounds.

We trained our model on the ADReSS Challenge dataset, a benchmark for AD speech analysis, and validated it on two independent corpora: the Delaware Mild Cognitive Impairment (MCI) corpus and a longitudinal dataset that we collected from President Ronald Reagan’s public speeches. This dual external validation provides strong evidence of generalizability across recording conditions, disease stages, and topical contexts.

To enhance clinical interpretability, we applied Explainable Artificial Intelligence (XAI) methods to identify which speech features most strongly influenced model decisions. By making the prediction process transparent, our approach supports clinician trust and facilitates the integration of speech biomarkers into clinical workflows.

From a translational perspective, our long-term goal is a patient-centered digital health tool that enables self-administered speech recordings via smartphones or online platforms for early AD screening and continuous monitoring. Such an approach could reduce diagnostic burden, improve accessibility, and empower older adults to engage proactively in their cognitive health management.

The remainder of this paper is structured as follows: Section 2 provides a review of relevant literature and highlights existing gaps in speech-based AD detection. Section 3 outlines the methodology and datasets used in this study. Section 4 explains the feature extraction process and examines how AD affects various speech characteristics. Section 5 presents the results along with model interpretation. Section 6 offers a discussion of the findings, their clinical implications, and the study’s limitations. Finally, Section 7 concludes the paper and discusses future directions.

## 2. Related Work

Extensive research has examined the impact of AD on language, with a significant focus on lexical features. Early evidence suggests that AD primarily affects lexical retrieval, which can manifest as word-finding difficulties and reduced vocabulary richness (Kurdi, 2024a; Mirheidari et al, 2018; Fraser, 2016; Garrard, 2005; Hebert et al, 2005; Hier et al., 1985). Clinically, lexical changes serve as early indicators of cognitive decline and have been used to support diagnosis in both research and practice. Other studies focused on the syntax of sentence (Nasiri et al, 2022; Eyigoz et al, 2020). Studies combining lexical and acoustic features have consistently reported improved AD detection, highlighting the potential of multimodal approaches (Cummins et al., 2020).

More recently, speech has emerged as a promising non-invasive biomarker for AD, offering scalable and accessible tools for early screening. Research has progressed along two main directions: some studies investigate theoretical correlations between AD and specific speech characteristics (Saeedi et al., 2024; Xue et al, 2021; Mueller et al., 2018; Ranasinghe et al, 2017; Yancheva et al, 2015; Forbes-McKay et al., 2013), while others focus on automated AI-based detection models (e.g. Kurdi, 2024b; Luz et al 2020; Jarrold et al, 2014). Both approaches aim to identify clinically relevant speech changes that can detect AD before significant cognitive decline occurs.

Phonetic and phonological speech features have also been explored. Early evidence suggested that phonological and articulatory impairments appear in the earliest stages of AD (Croot et al., 2000), and more recent studies confirm that these deficits persist throughout disease progression (Luchesi Cera et al., 2023). Features such as pause rate and segment duration have been linked to AD severity (Fernández, 2022; Cho et al., 2022). Clinically, these markers may indicate subtle disruptions in speech motor control and temporal organization, providing complementary information to lexical deficits. Remote speech assessments have also been shown to reliably detect amyloid-beta pathology in cognitively unimpaired individuals, reinforcing the potential of speech as an early, objective biomarker (Van den Berg et al., 2024).

Acoustic and prosodic features, such as speech rate, pitch variation, and temporal rhythm, have been extensively investigated. Studies report slower speech, increased pauses, and temporal disorganization in AD patients, observable across tasks like picture description or reading aloud (Hoffmann et al., 2010; López-de-Ipiña et al., 2013; Martínez-Sánchez et al., 2013; Satt et al., 2014). These features correlate with disease severity and can provide quantifiable indicators for clinical monitoring (Parlak et al., 2023).

Recent biomarker-linked speech studies indicate that early Alzheimer’s pathology can manifest as measurable disruptions in vocal control and temporal organization. Young et al. (2024) showed that speech-timing irregularities during memory recall tasks correlate with regional tau PET burden, suggesting that microprosodic and temporal instability may reflect early corticolimbic neurofibrillary degeneration. Complementarily, van den Berg et al. (2023) reported that naturalistic speech features, including pausing behavior, articulatory variability, and reduced fluency, track with cognitive vulnerability in preclinical cohorts and maintain reliability under remote acquisition, supporting their value as scalable digital biomarkers. The high-importance predictors in our model, phonatory instability (jitter and shimmer), temporal disruption, articulatory imprecision, and prosodic flattening, align with these pathology-linked feature classes and map onto degeneration of frontotemporal and temporoparietal speech–language networks.

Recent machine learning approaches highlight the diagnostic power of integrating lexical, syntactic, phonetic, and acoustic features. For example, Gonzalez-Atienza et al. (2021) demonstrated that mild AD is associated with shorter, slower, and more effortful speech, with increased disfluencies and pauses, achieving competitive classification performance. The ADReSS Challenge (Luz et al., 2020) has further established benchmarks for speech-based AD detection, with models leveraging fine-tuned language models (BERT, ERNIE) or combined audio-lexical features achieving accuracy about 89% (Yuan et al., 2020; Edwards et al., 2020; Balagopalan et al., 2020). Ensemble learning approaches integrating tree-based and neural classifiers have also improved robustness and interpretability (Pérez-Toro et al., 2022).

Despite these advances, systematic analyses of phonological and prosodic features remain limited. Most studies prioritize classification accuracy rather than exploring the underlying speech changes caused by AD. To our knowledge, no prior work has comprehensively examined prosodic, acoustic, and phonological speech structures from both empirical and conceptual perspectives. Our study addresses this gap by combining rigorous feature-based classification with in-depth linguistic analysis, aiming to provide clinically interpretable insights and explainable AI tools for early AD detection.

## 3. Methodology

Four ranking methods that yielded the highest classification performance were selected: ANOVA, Chi-Square (χ²), Information Gain, and ReliefF, along with Minimum Redundancy Maximum Relevance (mRMR). Feature ranking using the first four methods was performed with the Orange^1^ library, while mRMR ranking was carried out using the pymrmr Python library^2^ as part of the process of eliminating partially redundant features.

### 3.1 Objective

The primary objective of this paper is to systematically investigate the discriminatory effect of speech features in differentiating between patients with Alzheimer’s disease (AD) and healthy individuals. This exploration aims to provide a foundation for developing machine learning classifiers capable of performing this task automatically. The work is divided into two main phases: first, an individual analysis of how AD impacts a large number of speech features divided into three subgroups: phonological, acoustic, and prosodic features. Secondly, this study aims to identify the most effective combination of features and input length when necessary to optimize classification performance.

### 3.2 Hypotheses

As outlined in Section 2, several previous studies have shown that Alzheimer’s disease (AD) has a significant impact on speech. This leads us to hypothesize that speech in individuals experiencing healthy aging is less affected compared to those with AD. Moreover, based on the speech characteristics explored in earlier research, we expect AD to influence all the speech aspects examined in this study. We anticipate that AD will cause significant changes in prosody, acoustic features, phonetics, and phonology. Given that no comprehensive studies have systematically explored the full impact of AD on speech, it is difficult to predict which specific features will be most affected. Finally, considering AD’s impact on continuous acoustic and prosodic features, we hypothesize that it will also affect higher-level phonological features. Additionally, we propose that there may be a positive complementarity between the three feature subgroups, because they cover different aspects of speech, which could enhance AD classification results.

### 3.3 Datasets

The study uses the dataset provided by the ADDReSS challenge organizers (Luz *et al*., 2020), which is a subset of the DementiaBank corpus, part of the larger TalkBank project (MacWhinney et al., 2019). This dataset includes recordings and transcripts collected with the Cookie Theft picture description task, with 54 subjects in the training set and 24 subjects in the test set. The training set consists of 108 narrative samples, while the test set includes 48. Both speech audio files and transcripts were analyzed in this study. The ADDReSS dataset was chosen for its balanced representation of age, gender, and conditions, as described by Luz et al. (2020). Additionally, since the dataset has been used in several previous studies, it enables a comparison of our results with those of earlier research utilizing this dataset.

To evaluate the generalizability of our model beyond the ADReSS Challenge dataset, we conducted external validation using the **Delaware Corpus**, a recent addition to the DementiaBank repository. This corpus includes spontaneous speech samples from **individuals with Mild Cognitive Impairment (MCI)** and **cognitively healthy controls**, collected independently from ADReSS. Participants completed a range of **narrative and descriptive tasks**, including the *Cookie Theft* picture description, *Cinderella* story retelling, procedural instructions (e.g., “how to make a sandwich”), and personal narratives. Speech recordings are accompanied by transcriptions in CHAT format and metadata including cognitive assessments such as the **Montreal Cognitive Assessment (MoCA)** and **Boston Naming Test**. Only manually reviewed transcripts, free of automatic speech recognition (ASR) errors, were used in our evaluation to ensure data integrity. The Delaware dataset offers both linguistic and cognitive diversity and represents a clinically relevant benchmark for external validation. Its independent origin, varied task design, and distinct participant cohort make it a valuable resource for assessing model robustness in real-world diagnostic scenarios.

Previous studies (Wang, 2020; Berisha, 2015) have suggested that President Ronald Reagan showed early signs of Alzheimer’s disease during his second term, well before his official diagnosis in 1994. To examine this, forty five-minute passages were sampled from his public speeches. Twenty, five-minute passages, were taken from speeches delivered between 1964 and 1980, before any presumed symptoms, and twenty others were taken from speeches between 1988 and 1990, when signs of impairment were thought to emerge. All passages underwent **manual quality inspection** to ensure accuracy and consistency. Segments containing **multiple speakers**, **background noise**, **music**, **applause**, or other non-speech artifacts were excluded from the dataset. The speeches were retrieved from the Reagan Library Archives in MP4 format, converted to WAV using *pydub*, and transcribed with the *speech_recognition* library and the Google Speech-to-Text engine.

The ADDReSS challenge, the Delaware, and the Regan datasets used reflect real-world clinical populations with varying cognitive impairment levels, ensuring that the developed model is applicable across typical clinical scenarios.

As illustrated in Figure 1, three types of features were extracted: acoustic, prosodic, and phonological. Acoustic and prosodic features were derived directly from the audio wave files. Phonological features were obtained from transcriptions, including those provided by the ADReSS Challenge that were obtained manually and those that we generated via automatic speech recognition (ASR) to assess its impact. These feature sets will be used alone or in combination to train classifiers with different machine learning algorithms.

**Figure 1.**
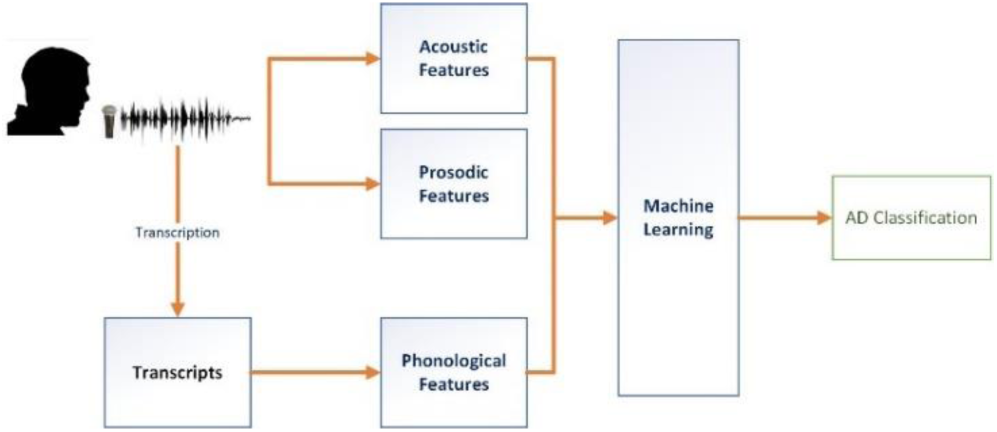
Outline of the work conducted in this paper

## 4. Features

Due to the extensive number of features examined in this study, a simplified ranking system was used to facilitate comparison. In this system, a ranking of A indicates that a feature is within the top 10%, B represents the top 25%, C the top 50%, D the top 75%, and a dash (-) signifies that the feature falls outside the top 75%.

To address redundancy, a two-stage feature selection was applied. First, features within each category (e.g., MFCCs) were ranked using mRMR, retaining the top 10% per category. As its names indicate, mRMR picks those features that reduce the redundancy, which is expected in this case given the high similarity between the features of the same type. Second, a global ranking was performed across the selected features to identify the most informative predictors.

### 4.1 Acoustic Features

Speech acoustics reflect cognitive and emotional states and are widely used in automatic speech recognition, speaker identification, and detection of mental health and neurodegenerative disorders (Deng et al., 2022; Edwards et al., 2020; Pérez-Toro et al., 2022). Acoustic features were extracted from all speech segments using openSMILE v2.1, a widely used open-source toolkit for automatic speech analysis, particularly in emotion and affective computing (Eyben et al., 2010, 2013), and extensively applied in Alzheimer’s disease research (Luz et al., 2020; Christensen et al., 2020; Pappagari et al., 2020; Edwards et al., 2020). In total, 6,056 acoustic features were generated (Table A1).

#### 4.1.1 Audio Spectral Features

Audio spectral features capture multiple characteristics of speech, including Relative Spectral Transform-Perceptual Linear Prediction (RASTA) and Relative Spectral Filtering. Each characteristic is quantified using standard statistical descriptors such as mean, standard deviation, quartiles, minimum, maximum, skewness, and kurtosis. Audio-based features have previously been applied to Alzheimer’s disease (AD) detection; for example, Shah et al. (2020) used spectral features within a classification framework and reported an accuracy of 60.19. Given the large number of spectral features available, only the top 10 features, based on ranking across the five applied feature-selection methods, are presented.

As shown in Table A2, statistical variations of the Rfilter features consistently ranked highest. All listed features appeared within the top 10% in at least three ranking methods, suggesting that these spectral characteristics are strongly influenced by AD.

#### 4.1.2 MFCC Features

Mel-Frequency Cepstral Coefficients (MFCCs) represent the short-term power spectrum of speech by applying a cosine transform to the logarithm of the power spectrum scaled on the mel frequency scale. MFCCs are widely used for feature extraction in speech and audio processing, including automatic speech recognition and speaker identification. Given their ability to capture spectral patterns associated with articulatory and phonatory changes, they have been extensively applied in speech-based Alzheimer’s disease (AD) detection. However, prior findings have been mixed. Martinc and Pollak (2020) reported an accuracy of 0.57 using MFCCs with an SVM classifier, whereas Balagopalan et al. (2020) found that MFCCs alone did not significantly distinguish AD from healthy controls. Cummins et al. (2020) used MFCCs within a Bag-of-Audio-Words framework, achieving an accuracy of 0.611 and an F1 score of 0.604.

In our analysis, the top 10 MFCC features (Table A3) were predominantly associated with the Simple Moving Average (SMA). Unlike the top-ranking audio spectral feature (audSpec_Rfilt_sma_de[21]_meanSegLen), none of the MFCC features appeared within the top 10% across all feature-ranking algorithms. This suggests that while MFCCs capture relevant acoustic markers of AD, their discriminative value is influenced by the choice of feature-selection method and is enhanced when combined with complementary acoustic and prosodic features.

#### 4.1.3 Pulse Code Modulation (PCM)

Pulse-code modulation (PCM) features capture basic signal energy and spectral properties of speech. Using openSMILE, we extracted three core PCM low-level descriptors: root mean square (RMS) energy, Fast Fourier Transform magnitude (FFT mag), and zero-crossing rate (ZCR). These features have previously shown sensitivity to speech changes in Alzheimer’s disease and related dementias (Kim et al., 2018; Shah et al., 2020; Lal et al., 2024).

After intra-group mRMR ranking, the top 10 PCM features were retained (Table A4). Notably, no RMS-based features ranked in the top 10; all selected features were derived from FFT magnitude, indicating that spectral energy distribution carries greater discriminative information than overall signal energy in this older-adult cohort.

#### 4.1.4 Log Harmonic to noise and Voice Final Unclipped

The harmonics-to-noise ratio (HNR) quantifies the relative energy of periodic versus aperiodic components in speech, with lower values indicating increased vocal irregularity commonly observed in aging and neurodegenerative conditions (Martínez-Nicolás, 2020). The logarithmic version (Log HNR) is used for better numerical stability. Although one study found no significant group differences in HNR between individuals with Alzheimer’s disease (AD) and healthy controls (Parlak et al., 2023), Log HNR-derived features have contributed to successful speech-based AD classification in multiple prior works (Vigo et al., 2022).

We also included voicing-related features from final unclipped speech segments (voicing probability at phrase endings), which influence perceived speech naturalness and have been employed in automatic speech recognition and clinical voice assessment (Catts & Jensen, 1983).

The top six Log HNR and voicing features retained after intra-group ranking are listed in Table A5. As shown in Figure B1 and Table 5, these features consistently ranked lowest across selection methods, partly reflecting the limited number of variants in this functional group. In contrast, audio spectral features (blue) achieved the highest rankings, benefiting from greater variant diversity and stronger discriminative performance in this older-adult cohort (Table A2).

Additionally, Figure B2 shows that the log HNR and voicing features consistently fall into the lowest subgroup, which is consistent with the low rankings seen in Table A5. This can be partly explained by the smaller number of variants in this subgroup compared to others. In contrast, the Audio Spectral Features, represented in blue, consistently achieve the highest rankings across all feature selection methods. This strong performance is likely attributed to the larger number of variants in this subgroup, as well as the higher number of features that made it into the top ranks, as indicated in Table A2.

### 4.2 Prosodic Features

Cognitive impairments, particularly in memory and executive function, have been linked to altered prosodic patterns, including reduced pitch-related measures and increased jitter, shimmer, and amplitude variability (Martínez-Sánchez et al., 2013; Meilán et al., 2012). Prosody reflects rhythm, melody, and intonation in speech and contributes to meaning, emotion, and emphasis. It can be broadly categorized into prosodic acoustic features and prosodic phonological features. Prosodic acoustic features capture continuous aspects of speech, such as fundamental frequency (F0) variation, speech rate, and pitch; these are examined in this section and summarized in Table A6. Prosodic phonological features, which describe discrete elements such as syllables and phoneme segments, are addressed in Section 4.3.

A total of 470 prosodic features were extracted. Because jitter and shimmer include a large number of highly redundant measures, a two-stage feature selection process was applied. First, the most informative jitter and shimmer features were identified; the top 10 were then combined with the remaining prosodic feature sets.

#### 4.2.1 Fundamental Frequency Variations

Fundamental frequency (F0), perceived as pitch, is the lowest frequency of a periodic speech signal and reflects the vibration rate of the vocal folds. Variations in F0 capture changes in pitch and intonation over time. This section focuses on overall statistical patterns of F0 variation, including mean, standard deviation, linear regression indicators, and quantile-based measures. In total, 39 F0 variation features were extracted using the openSMILE library.

Previous studies have linked F0 characteristics to cognitive decline, although results remain inconsistent. Parlak et al. (2023) reported a significant negative correlation between Mini-Mental State Examination (MMSE) scores and F0. Meilán et al. (2012) found no association between Alzheimer’s disease (AD) and mean F0, whereas Bae et al. (2023) observed strong correlations between AD and both mean F0 and the 20th percentile of F0. Similarly, Huang et al. (2024) identified significant associations for five of seven F0 measures examined. Given the extensive number of F0 features, only the top 10 ranked measures are presented.

As shown in Table A7, three variants of the Linear Predictive Coefficient (LPC) were among the top four F0 features; however, due to redundancy among these measures, their ReliefF scores were comparatively lower. Other highly ranked F0 descriptors included interquartile ranges, minimum and maximum values, regression-based measures, and rise time.

#### 4.2.2 Pitch

Speech pitch is the perceptual attribute that conveys how high or low a voice sounds and is determined by the frequency of vocal fold vibration. Pitch plays a central role in expressing intonation, emotion, and emphasis in spoken language. Not all speech sounds are voiced, however; for example, the sounds *[s]* and *[p]* are voiceless because they do not involve vocal fold vibration (Kurdi, 2017). Speakers also rapidly adjust their vocal output to compensate for brief perturbations in pitch within their auditory feedback (Ranasinghe et al., 2017). Pitch-related features have been used in several studies to detect Alzheimer’s disease (AD), with promising results reported by Huang et al. (2024), Ambrosini et al. (2019), and Yu et al. (2015).

As shown in Table A8, two pitch features, meanP and pearModeFirstP, ranked highest across all four feature ranking methods. Other features, such as iqrP, skewP, kurtFisherP, and kurtPearsonP, received high rankings from three methods but a lower ranking with relifF, possibly due to partial redundancies among these features.

#### 4.2.3 Amplitude

Amplitude is a key prosodic acoustic parameter reflecting the loudness of speech, defined as the maximum displacement of vibrating particles from their resting position. Prior research shows that individuals with Mild Cognitive Impairment (MCI) exhibit weaker vocal intensity than healthy older adults, characterized by increased dysphonia and a higher ratio between the first harmonic amplitude and the F3 amplitude during phonation (Themistocleous et al., 2020). Similar amplitude-related abnormalities have been reported in individuals with Alzheimer’s disease (AD) compared with healthy controls (Tanaka et al., 2017). Amplitude-based features have also been incorporated into recent AD classification studies using acoustic markers (Luz et al., 2020; Hason & Krishnan, 2022). The top 10 ranked amplitude features are summarized in Table A9.

As shown in Table A9, the feature-ranking methods displayed greater disagreement for amplitude features than for other acoustic subtypes. For example, mean amplitude received an A ranking with information gain but a D ranking across the remaining methods. Some features, such as the interquartile range (IQR), consistently received A or B rankings, whereas others, including variance, mode, delta, and Pearson Mode Second, ranked consistently low across all ranking approaches.

#### 4.2.4 Intonation

Intonation refers to the melodic contour of speech, reflected through pitch variations over time. Several studies have investigated how Alzheimer’s disease (AD) alters intonational patterns (Mueller et al., 2018). For example, Forbes-McKay et al. (2013) compared melodic contours in approximately 60 participants and found that individuals with mild to moderate AD exhibited reduced intonational variation during both simple and complex description tasks. Intonation features have also been incorporated into AD speech-classification models (König et al., 2015). The top 10 intonation features identified in our analysis are shown in Table A10.

Across feature-ranking methods, substantial variability was observed. However, the *Ratio peaks* feature consistently ranked among the highest, whereas the *Mean* feature consistently appeared near the bottom across all ranking methods.

#### 4.2.5 Articulation Rate and Patterns of Unfilled Pauses

Articulation rate, the speed of producing linguistic units, varies with emotional state, context, speaker characteristics, and language. Slower speech is linked to word-retrieval difficulties (Satt et al., 2014; Pu & Zhang, 2025) and, along with speech tempo and hesitation ratio, distinguishes individuals with Alzheimer’s disease (AD) from healthy controls (Hoffmann et al., 2010; Huang & Yang, 2022). Speech rate is among the most discriminative features for differentiating AD from healthy aging (Martínez-Sánchez et al., 2013), while pause patterns add further diagnostic value (López-de-Ipiña et al., 2013). Changes in speech duration and rhythm may signal early AD (Meilán et al., 2012), and slower speech is also reported in other dementias, including Parkinson’s disease (Skodda & Schlegel, 2008). Recent studies have successfully incorporated speech rate and tempo into AD classification (Huang et al., 2024).

We analyzed three speech-rate features, ratioWords, ratioChars, and ratioSylls, defined as the number of linguistic units divided by total utterance duration (Table A11). Despite prior evidence linking slower speech to AD, these features consistently ranked low across all four feature-selection methods, with syllables per second performing worst.

Pauses, while partly driven by respiration, also convey linguistic and speaker-specific information (Kurdi, 2002). AD speech exhibits longer, more frequent, and more variable pauses (Pastoriza-Domínguez et al., 2022; Hason & Krishnan, 2022; Martínez-Nicolás et al., 2020), making silence a widely used marker of cognitive impairment that reflects delays in speech planning and production (Gonzalez-Atienza et al., 2021; Pappagari et al., 2020; Yuan et al., 2020). However, there is no consensus on the minimum pause duration, with cutoffs ranging from 10 ms (Skodda & Schlegel, 2008) to 50 ms (Goberman et al., 2005) or 200 ms (Bunton & Keintz, 2008; Gravelin & Whitfield, 2017).

We computed the ratio of silent segments to total signal duration using thresholds of 10, 25, 50, 100, and 200 ms. Detection was challenged by overlapping speech and background noise. As shown in Table A12, none of the silence features ranked highest across methods, though the shortest threshold (10 ms) performed best. No consistent pattern emerged across pause durations, as longer thresholds sometimes ranked higher than shorter ones and vice versa.

#### 4.2.6 Jitter and Shimmer

Jitter and shimmer represent micro-prosodic fluctuations in pitch and amplitude, respectively, and are commonly used to assess voice quality and detect vocal pathologies (Wertzner et al., 2005; Teixeira et al., 2013). Recent findings indicate that individuals with Alzheimer’s disease (AD) exhibit significantly higher jitter and shimmer values compared with healthy controls (Parlak et al., 2023). These measures have also been incorporated into automated AD classification studies (Hason & Krishnan, 2022). The top 10 shimmer and jitter features identified in our analysis are presented in Tables A13 and A14.

As shown in Tables A13 and 14, the rankings of similar features, such as jitterLocal_sma_range and shimmerLocal_sma_qregerrQ, vary considerably across feature-selection methods. At the same time, clear distinctions emerge between the highest- and lowest-ranked jitter and shimmer features, indicating substantial variability in their discriminative value.

#### 4.2.7 Comparison of the Prosodic Features by types

A comparison of the prosodic features is provided in Figure B3. Given that F0 features represent the largest group in the dataset, it is reasonable to expect them to dominate the ranked feature lists. As shown in Figure 2, this expectation is largely confirmed: F0 features account for 65.4% of the top features in the ANOVA ranking and 50% in the χ² ranking. Their prominence, however, decreases substantially under information gain (34.6%) and even more under ReliefF (7.7%). This reduction is consistent with the ReliefF algorithm’s tendency to penalize partially redundant features.

**Figure 2.**
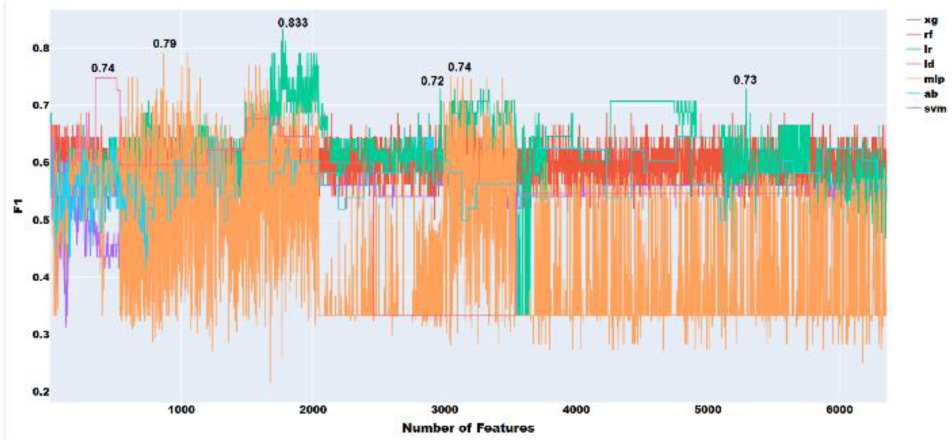
Feature selection experiments with all the acoustic features

Amplitude features exhibit the opposite pattern. They play a minimal role in the ANOVA and χ² rankings (3.8% and 7.7%, respectively) but appear more frequently in the information gain and ReliefF lists (11.5% and 26.9%, respectively). Pitch features, by contrast, maintain a relatively strong presence across all four ranking methods. Unfilled pause features do not appear among the top features in the ANOVA results and occupy only a minor share 3.8% in the χ², information gain, and ReliefF rankings.

### 4.3 Phonological Features

Phonology examines the formal structure of speech sounds, including segmental elements (e.g., phonemes, syllables) and suprasegmental features such as intonation and stress. Several studies have explored phonological changes in Alzheimer’s disease (AD). Croot et al. (2000) reported early phonological and articulatory impairments in AD, and Luchesi Cera et al. (2023) found phonological disorders to be the most prevalent across mild, moderate, and severe AD stages compared with healthy controls.

Guided by prior work, including the Index of Phonetic Complexity (IPC) (Akielski, 2022), we analyzed nine features reflecting word-level phonemic structure, including grapheme count (nbLett), phoneme count (nbPhon), and the presence of specific consonant classes (labials, stops, nasals, dorsals, glides, fricatives, affricates, liquids). Features were organized into three subcategories: consonants, vowels, and supraphonemic features, with consonants further analyzed by manner and place of articulation (Kurdi, 2017).

For each utterance, we computed the ratio of each consonant or vowel type to the total number of phonemes, providing the phonological features used in the analysis.

#### 4.3.1 Consonant Features

Consonants are speech sounds produced by obstructing airflow at specific points in the vocal tract, defined as the place of articulation. In this study, we examined standard phonological categories, including bilabial, labiodental, dental, alveolar, post-alveolar, palatal, glottal, lateral, and coronal consonants, analyzing both place and manner of articulation. Consonant Place of Articulation (CPA) was represented by 18 features capturing distinct articulatory locations.

Dental consonants (/θ/, /ð/) ranked highly across all four feature-selection methods (Table A12), despite low overall frequency in English (/θ/: 37th of 39; /ð/: 11th). Their prominence in Alzheimer’s disease (AD) speech likely reflects compensatory lexical behavior, as individuals with AD overuse high-frequency function words such as “the,” “this,” and “that,” consistent with prior evidence that AD speakers rely more on closed-class words when semantic retrieval is impaired (Kurdi, 2024a). In contrast, bilabial, coronal, and lateral consonants consistently ranked low, suggesting these phonemes are less informative for distinguishing AD from healthy speech. Mid-ranked palatals showed no clear association with lexical patterns. These results indicate that specific consonant classes, particularly dentals, may serve as indirect markers of lexical-semantic deficits, supporting the hypothesis that AD affects subtle phonological patterns detectable via automated analysis.

Consonant Manner of Articulation (CMA), which reflects how airflow is modified during production, was represented by 27 features. Fricatives and fricative–affricate–liquid (friAffLiq) features consistently ranked highly (Table 13). Their prominence cannot be explained solely by frequency, as some fricatives (e.g., /ʃ/, /ʒ/) are rare, while others (e.g., /t/, /s/) are common, suggesting overlap with dental phonemes and reinforcing the relevance of specific consonant classes to AD-related speech changes.

#### 4.3.2 Vowel Features

In phonology, vowels are speech sounds produced without significant constriction in the vocal tract, allowing airflow to pass freely and typically forming the nucleus of syllables. Two ratios were computed for each vowel: the proportion of a specific vowel type to the total number of phonemes in a word (e.g., high-to-phon) and the proportion relative to the total number of vowels in the word (e.g., high-to-vow). Eight vowel types were analyzed as features (Table A14). Several features, including tense, syllabic, and round vowels, ranked highly across feature-selection methods (Table 14).

### 4.4 Misc. Features

This subset includes syllabic consonants, which can serve as the nucleus of a syllable, and diphthongs, complex vowel sounds that glide from one vowel to another within a single syllable. Table A19 presents the ratios of each sound type relative to consonants and to all phonemes, showing variable rankings. Overall, some features, such as syllabic consonants relative to consonants, ranked above average, whereas others, like diphthongs relative to consonants, ranked lower.

### 4.5 Supra-phonemic Structure

Supra-phonemic structures are key indicators of phonological complexity. These include general phoneme sequence features, such as consonant clusters (consCluster), defined as two or more consecutive consonants without an intervening vowel, and word-final consonants (endConson). Syllable structure also reflects phonological complexity, comprising an onset (consonants before the nucleus), a nucleus (typically a vowel or diphthong), and an optional coda (consonants after the nucleus). Two linguistic approaches measure syllabic complexity: categorical syllable complexity (Maddieson, 2007) and the sum of maximal onset and coda (Gordon, 2016; Easterday, 2019), which consider the sizes and types of phonemes in these positions. Accordingly, this study uses lenOnset and lenCoda, as well as vowel-type ratios within the syllable nucleus, back vowels (backSyll), front vowels (frontSyll), and diphthongal nuclei (diphSyll), and the average number of syllables per word (avgSyll).

Two features capture suprasegmental structure: primary stress (primStress), marking the strongest syllable, and secondary stress (secondStress), marking strong but less prominent syllables. Phonemic sequences were obtained from the CMU Pronouncing Dictionary; grapheme sequences were used when dictionary entries were unavailable.

As shown in Table A16, avgSyll and consCluster rank within the top 10% of features, demonstrating strong discriminative value and supporting the importance of deeper linguistic analysis. Coda consistently ranks higher than Onset, reflecting the optional nature of onsets in English. Stress-related features (primStress, secondStress) consistently rank low, indicating a limited association with Alzheimer’s Disease (AD).

Figure 3 illustrates that Consonant Manner of Articulation (CMA) features dominate rankings across all four feature-selection methods. Miscellaneous features consistently rank lowest. Under ReliefF, vowel and Consonant Place of Articulation (CPA) features receive lower rankings, whereas supra-phonemic features achieve relatively higher positions.

**Figure 3.**
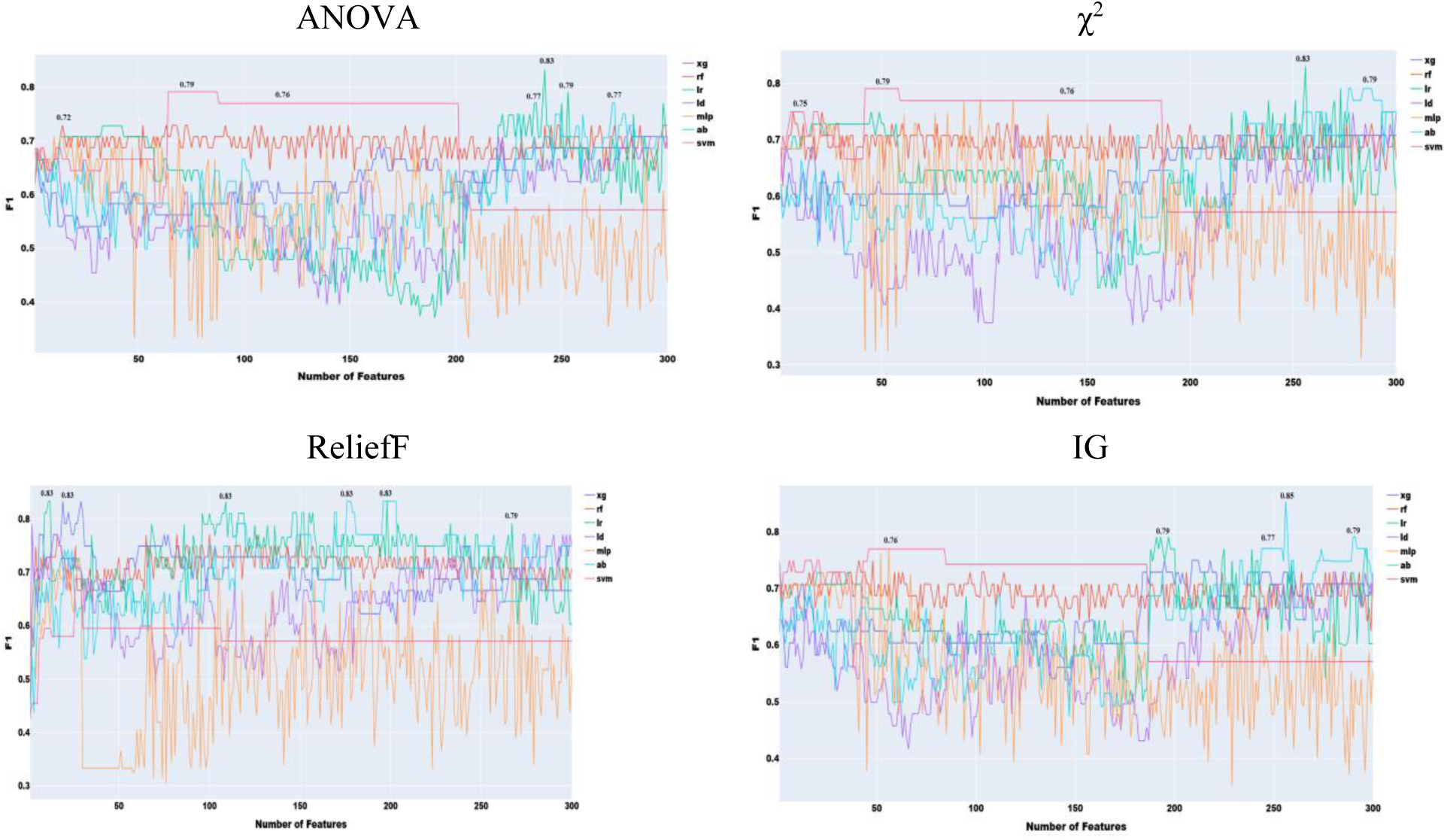
Feature selection experiments with acoustic features only (simple classifiers)

**Figure 4.**
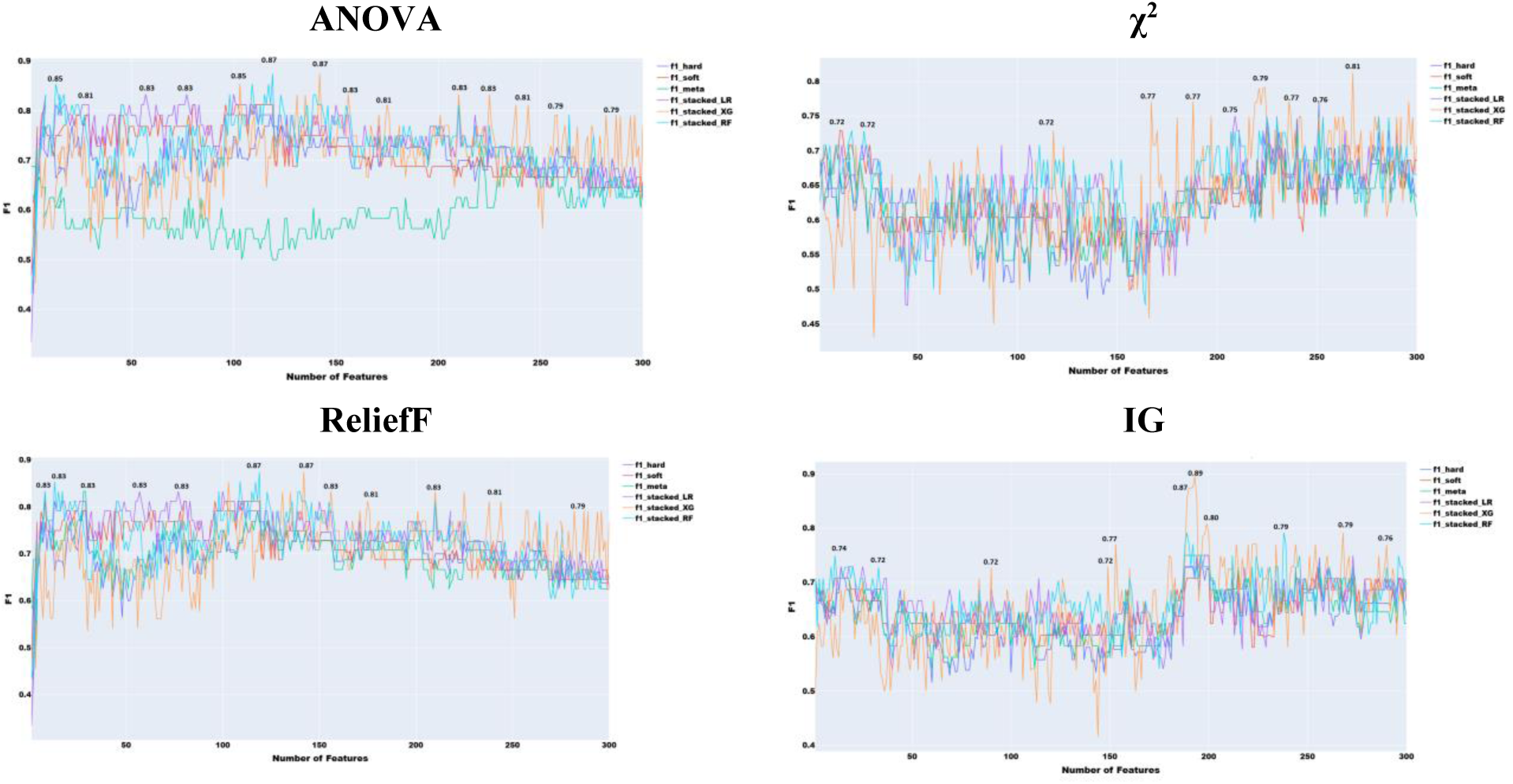
Feature selection experiments with acoustic features only (ensemble learning)

## 5. Results and Discussions

This section evaluates the predictive power of the speech features described in Section 4. The following machine learning (ML) algorithms were applied: Support Vector Machine (SVM), Logistic Regression (LR), Linear Discriminant (LD), Adaptive Boosting (AB) with 100 estimators, Bagging (Breiman, 1994),

Random Forest (RF) with maximum depth 2 (Ho, 1995), eXtreme Gradient Boosting (XG) (Chen & Guestrin, 2016a, 2016b) with parameters learning_rate=0.001, n_estimators=5600, max_depth=5, min_child_weight=1, gamma=0, subsample=0.8, colsample_bytree=0.9, objective=’binary:logistic’, seed=25, and a Multilayer Perceptron (MLP) with max_iter=200, hidden_layer_sizes=50, activation=’tanh’, solver=’adam’, and alpha=1e-8. Parameters were selected empirically to optimize performance. These algorithms were chosen based on preliminary experiments comparing them with alternatives such as Decision Trees and Naïve Bayes. All experiments were implemented in Python using Scikit-learn (Pedregosa et al., 2011) and XGBoost. Performance was evaluated using accuracy, recall, precision, and F1-score (Manning et al., 2008).

Ensemble learning, which combines multiple classifiers, was also explored given its successful application in AD classification (Sarawgi, 2020). Three ensemble methods were employed: hard voting, soft voting, and stacking. In stacking, a meta-classifier assigns optimal weights to each base classifier; here, Adaptive Boosting was used to learn these weights from base classifier probabilities. Ensemble results are reported only when they outperform individual classifiers.

Only F1-scores are presented, as other metrics offered no additional insight. F1 was selected both for its widespread use in ML experiments and to align with the ADReSS Challenge, where it is a formally adopted evaluation measure.

### 5.1 Experiments with Acoustic Features

To choose the most effective set of acoustic features and evaluate the performance of classifiers using only these features, we adopted the following approach as with the acoustic features. We progressively added features ranked by the four feature ranking methods, starting with the highest-ranked feature and moving eventually to the lowest. The results of this feature selection process are shown in Figure 5, with all the 6376 acoustic features and ANOVA feature selection.

**Figure 5.**
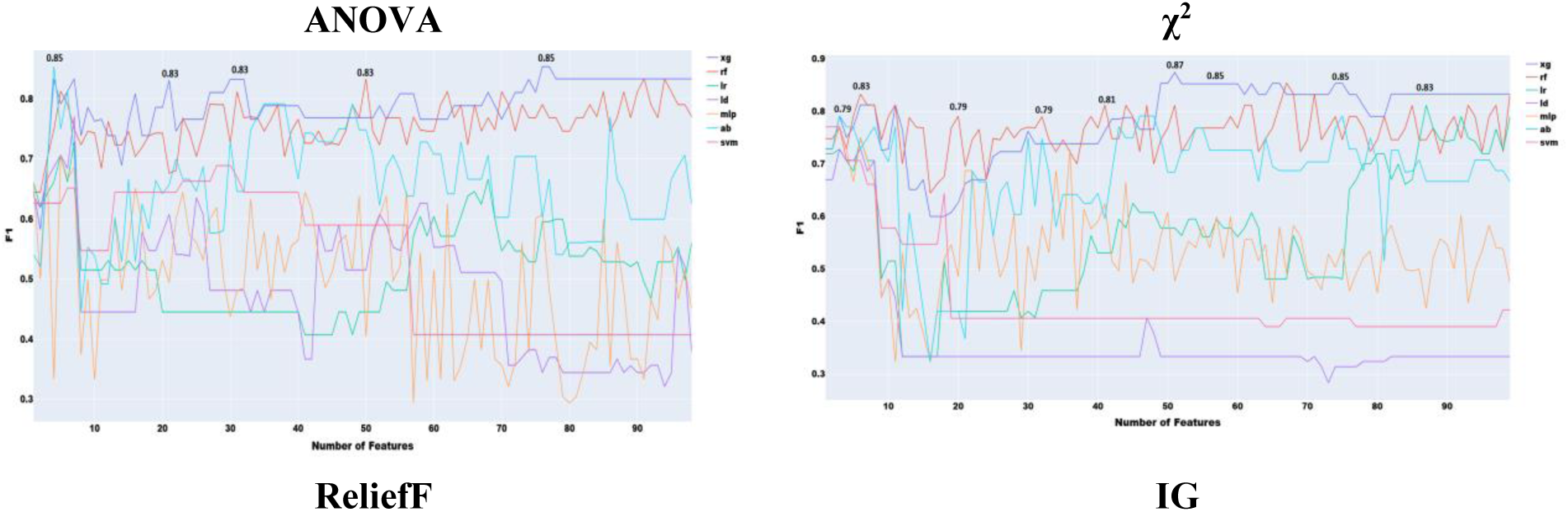
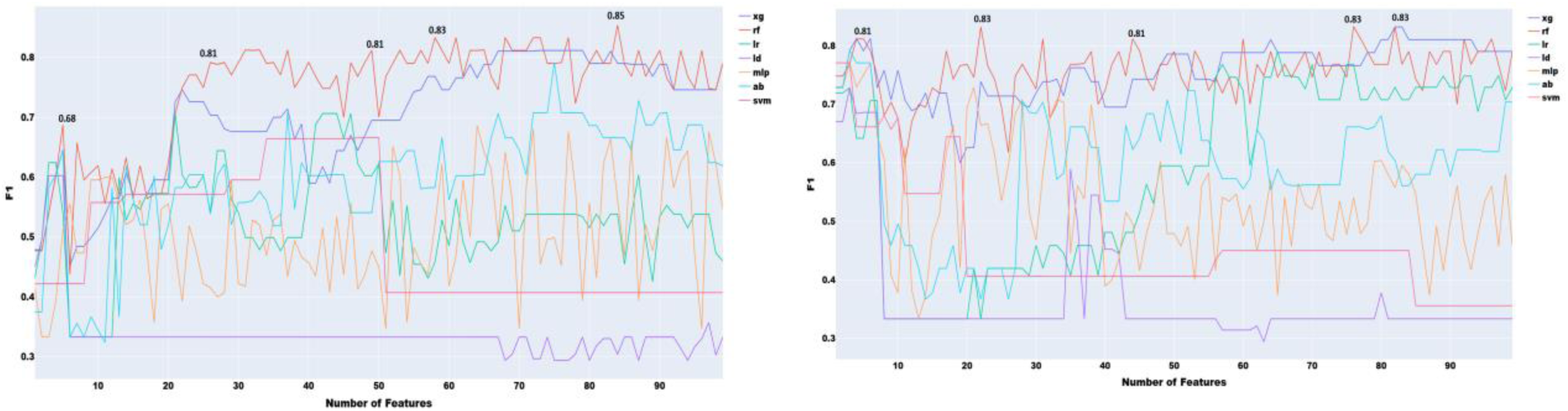
Feature selection experiments with prosodic features only (simple classifiers)

As shown in Figure 5, the highest performance using all acoustic features was an F1 score of 0.83 with logistic regression. To reduce redundancy in the large feature set, where many features capture different statistical descriptors of the same acoustic properties (e.g., MFCC, F0, jitter), we implemented a two-stage feature selection. First, mRMR was applied to extract the 300 most informative features. Second, this subset was ranked using four additional feature selection methods to identify the most effective combination (Figure 6).

**Figure 6.**
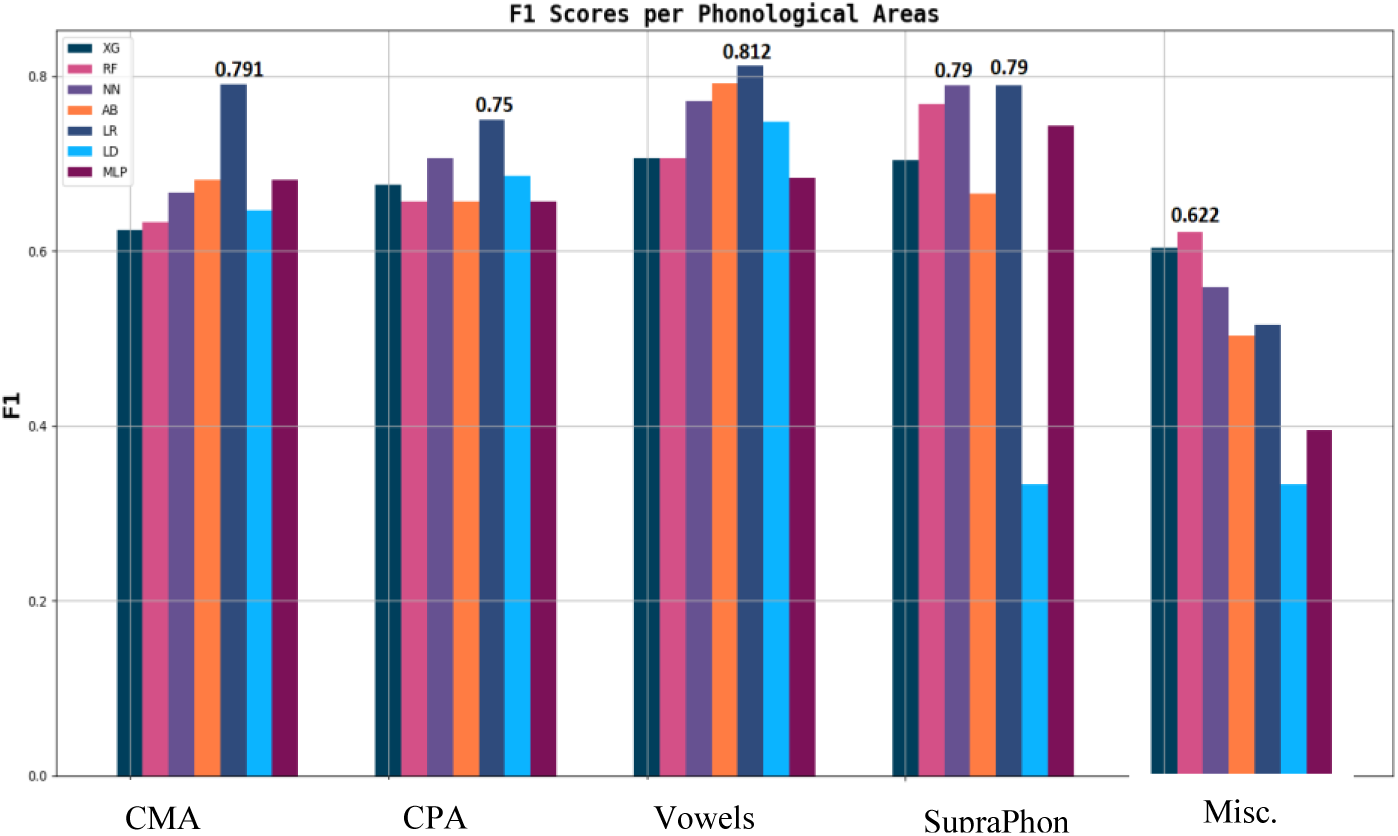
F1 scores of Phonological Areas

As shown in Figure 6, the highest performance with acoustic features was obtained using the information gain method, yielding an F1 score of 0.85. The remaining feature selection techniques reached their peak performance at an F1 score of 0.83. It is also notable that, with the exception of ReliefF, all methods required a relatively large number of features to achieve optimal results.

To further analyze the generalization behavior of our models, we inspected the confusion matrix of the best-performing classifier, adaptive boosting trained on the top 256 information gain features, which achieved the highest F1 score of 0.85.

As shown in Table A21, the classifier has similar rates of true negatives and true positives, suggesting a balanced error pattern. In other words, it performs equally well on both positive AD cases and control cases.

Ensemble learning experiments were conducted. Instead of using all the ML algorithms that we experimented with so far, only those that displayed a steady performance in Figure 5 were retained. Hence, we have used the following ML algorithm in the ensemble experiments: SVM, Random Forest, Adaptive Boosting, Logistic Regression, and XG Boost.

As illustrated in Figure 6, ensemble learning improved the F1 score by 4% when using acoustic features, achieving a score of 0.87 with ReliefF and stacked ensemble approach with both XGBoost and Random Forest. A score of 0.89 was achieved with Information Gain features selection combined with XGBoost with stacked ensemble approach.

### 5.2 Experiments with Prosodic Features

To select the best prosodic features and assess the performance of the classifiers with these features only, we operated in a similar way with the acoustic features. The results of these feature selection methods are presented in Figure 7.

**Figure 7.**
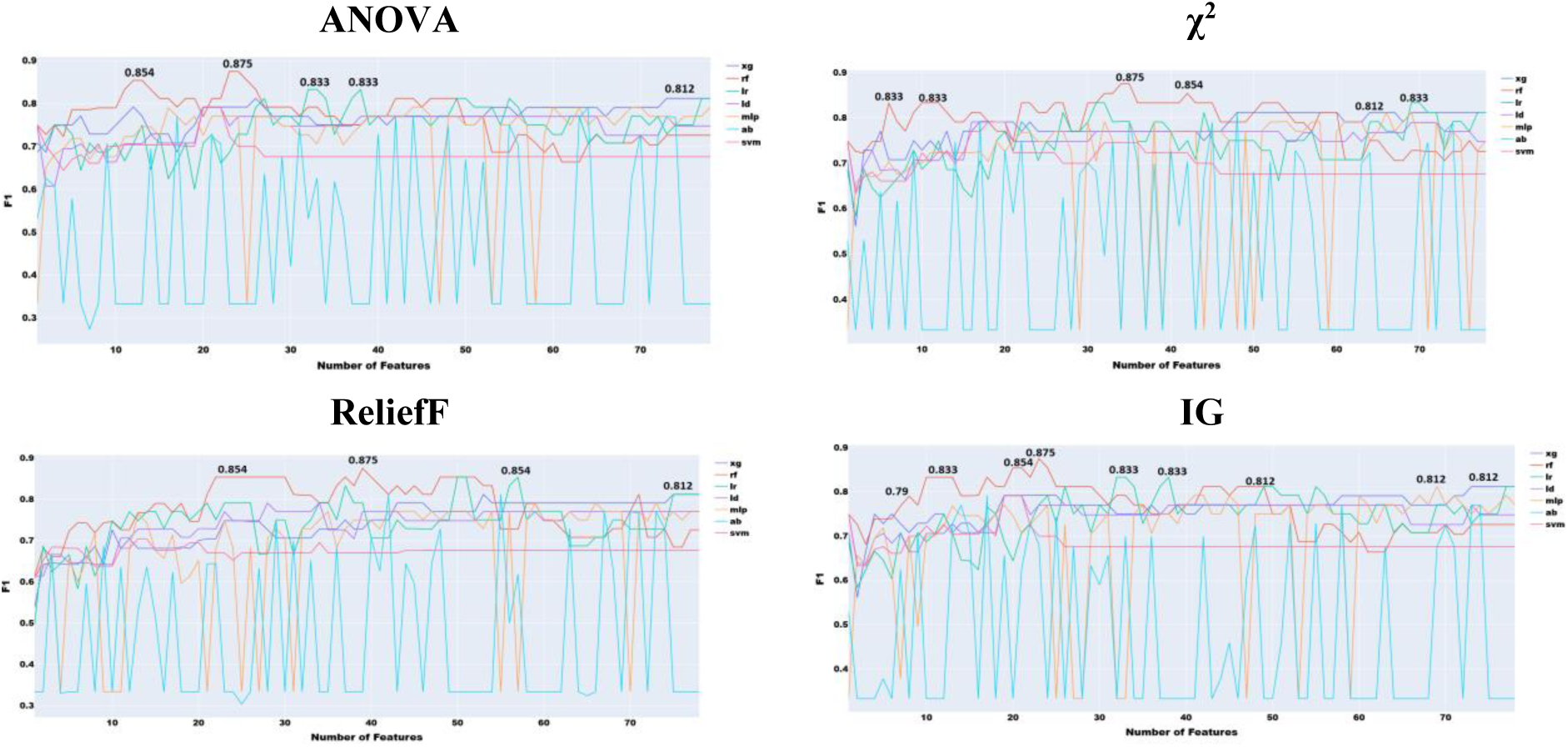
Feature selection experiments with phonological features only (human transcribed)

**Figure 8.**
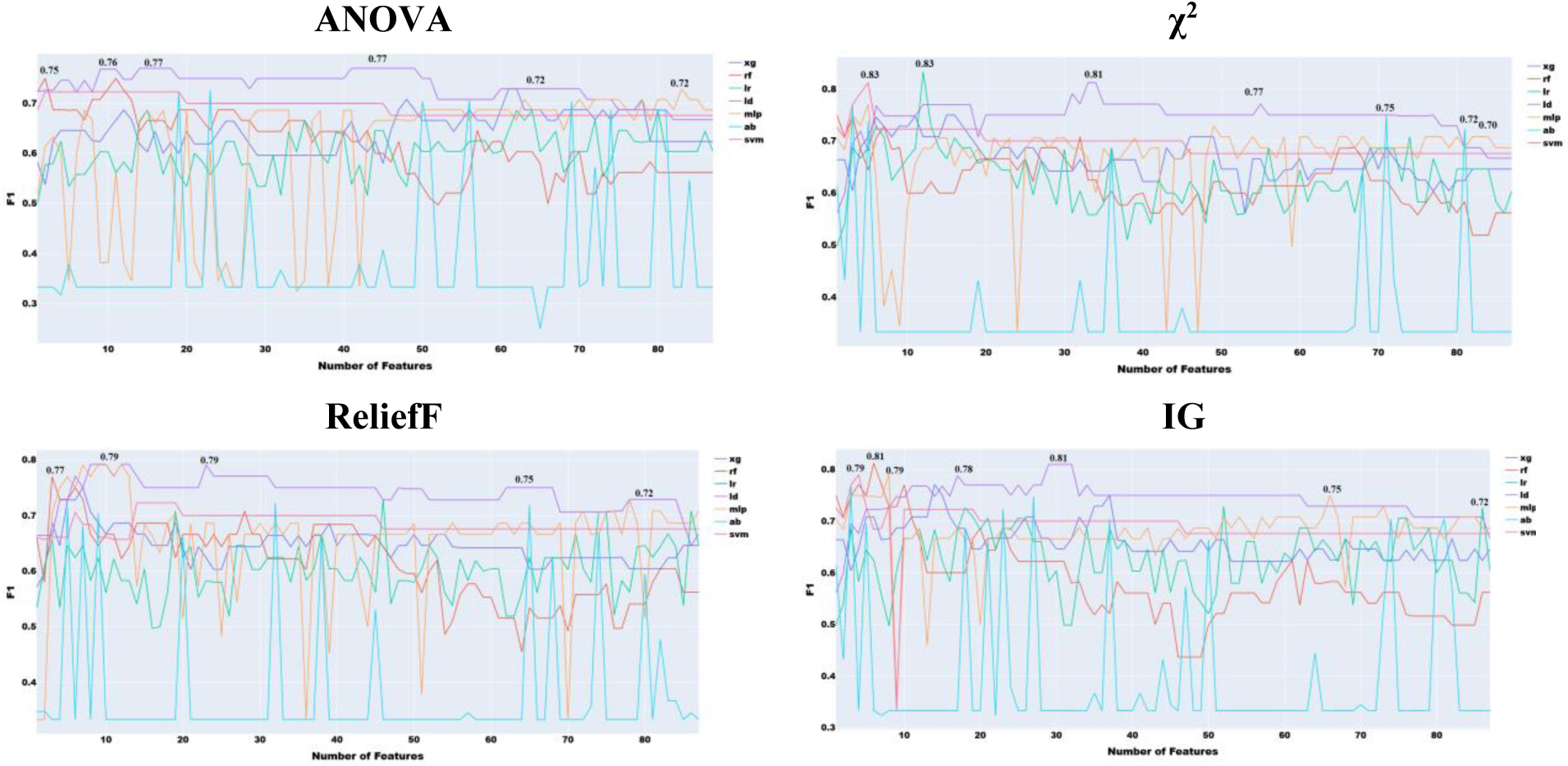
Feature selection experiments with phonological features only (ASR transcribed)

As seen in Figure 5, the best performance of 0.87 with prosodic features was yielded by χ^2^ with XG Boost.

The confusion matrix for the two classifiers that produced the best classification performance using the prosodic features and ANOVA, Adaptive Boosting with 4 features and XGBoost with 88 features, is shown in Table A6 to identify the error patterns.

As illustrated in Table A18, XGBoost combined with both χ^2^ and ANOVA demonstrates superior performance in detecting true positives and minimizing false negatives. However, this comes at the cost of increased false positives and a reduction in true negatives. Since false negatives are deemed more critical than false positives due to their potential to delay treatment, it can be concluded that XGBoost outperforms AD in this context.

### 5.3 Experiments with Phonological Features

#### 5.3.1 Evaluation of Phonological Areas

Organizing the features according to phonological domain or category enables a clearer examination of how AD affects each group of features relative to their linguistic functions. Methodologically, although the study does not aim to exhaustively capture every possible phonological feature, it includes a sufficiently broad set to allow the comparison to reflect overall patterns or trends.

A few key observations emerge from Figure 6. With the exception of the miscellaneous category, all phonologic feature types perform reasonably well, though their performance remains below the best results obtained using prosodic and acoustic features (Figures 5, 6 and 7, respectively). Vowel features yield the strongest classification performance, achieving an F1 score of 0.81 with logistic regression, which aligns with their high impact shown in Table 18 (Rankings of the vowels features). The number of features within each category does not appear to systematically influence performance. For example, although CMA includes nearly three times as many features as the vowel category, its results are only slightly lower, likely due to higher internal redundancy. Overall, the logistic regression model consistently delivers the best performance across all four linguistic types.

#### 5.3.2 Feature Selection Experiments

To identify the optimal subset of features across all linguistic types, we applied the four adopted feature selection methods. Each method ranks features differently, so we constructed a progressively expanding list, starting with the top-ranked feature and adding others incrementally until all features were included.

F1 results (Figure 11) indicate that adding more features does not always improve classification, likely due to internal redundancies. All four methods achieved the same peak F1 of 0.87, with Information Gain and ANOVA reaching optimal performance using fewer features. Despite the small training set, this suggests that phonetic and phonological features alone can yield strong classification performance.

Among machine learning algorithms, random forest (RF) consistently performed best across all feature selection methods, followed by XGBoost and logistic regression (LR). These findings underscore the importance of RF and LR, as also observed in the linguistic-type evaluation (Figure 6). In contrast, prior studies focusing on lexical features (e.g., Kurdi, 2024a) did not find a clear algorithmic advantage, highlighting that algorithm performance depends on the selected feature set.

As shown in Figure 9, all four feature selection methods achieved the highest F1 score of 0.87, comparable to top performances in the ADReSS Challenge, where models leveraged multiple linguistic levels. Achieving a similar score using only phonological features, without lexical or acoustic cues, suggests that phonological characteristics alone are highly informative for Alzheimer’s disease (AD) diagnosis, highlighting the potential of phonology-based models for early and accurate detection.

**Figure 9.**
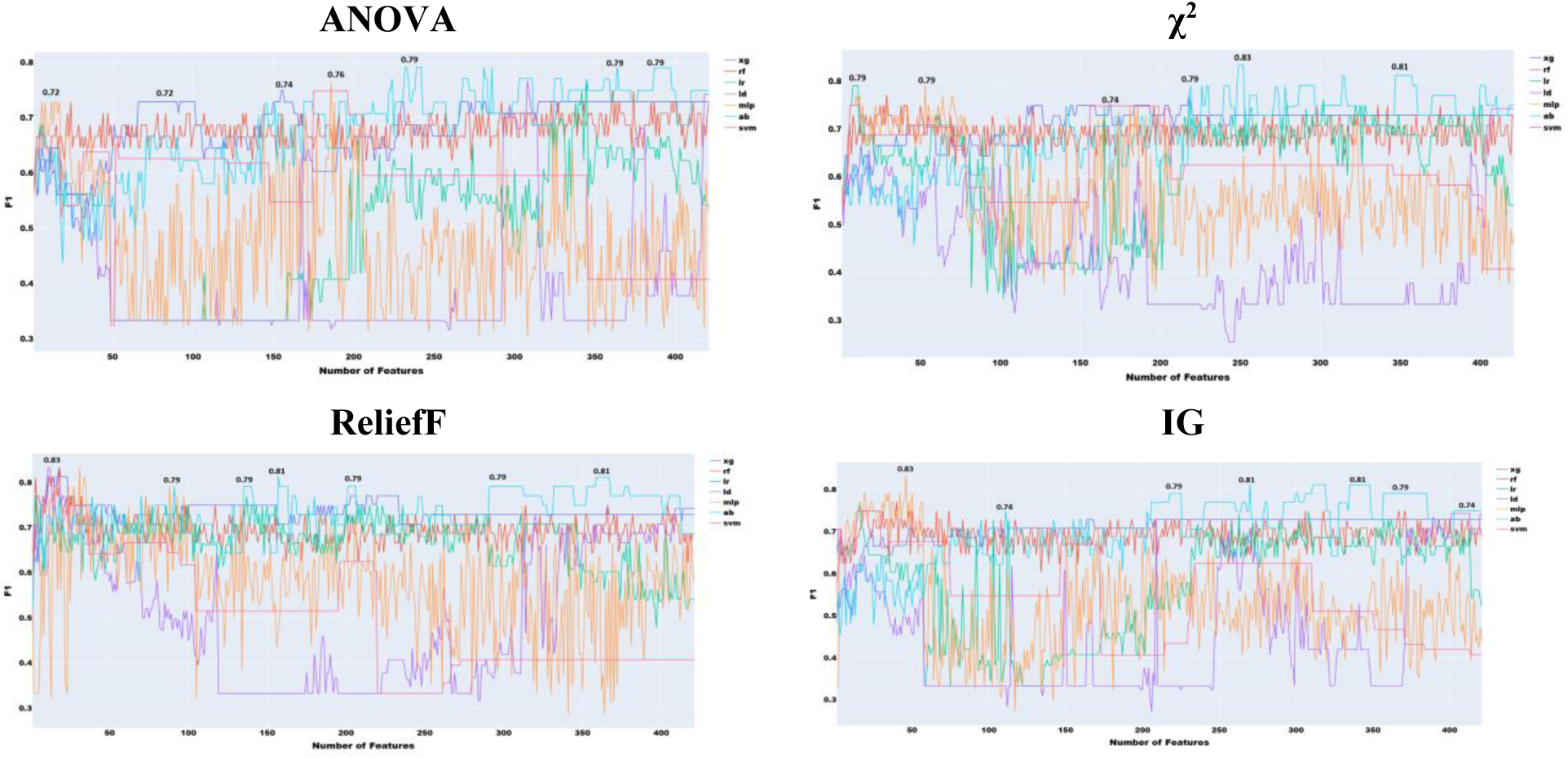
Feature selection experiments with prosodic and acoustic features combined

Error patterns are detailed in Table 23, which presents confusion matrices for the classifier using 23 ANOVA-selected phonological features with XGBoost. The equal number of false positives and false negatives indicates balanced performance across subgroups.

Prior experiments used manually transcribed speech. To assess the impact of automatic speech recognition (ASR), we tested a hybrid approach combining Google and Sphinx modules from Python’s *speech_recognition* library. The Google engine was prioritized, with Sphinx as backup. Results are reported in Figure 10.

**Figure 10.**
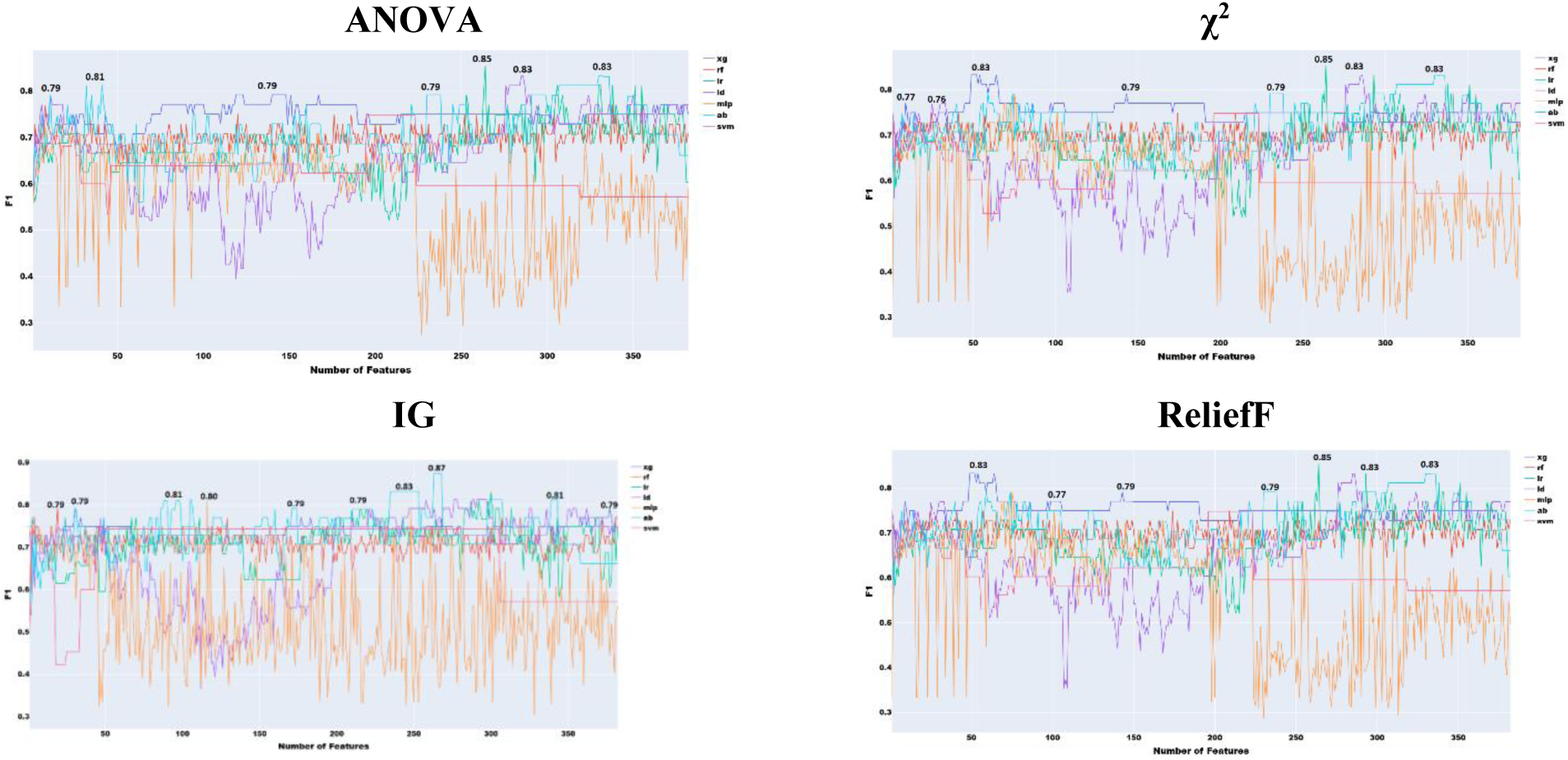
Feature selection experiments with acoustic and phonological features

**Figure 11.**
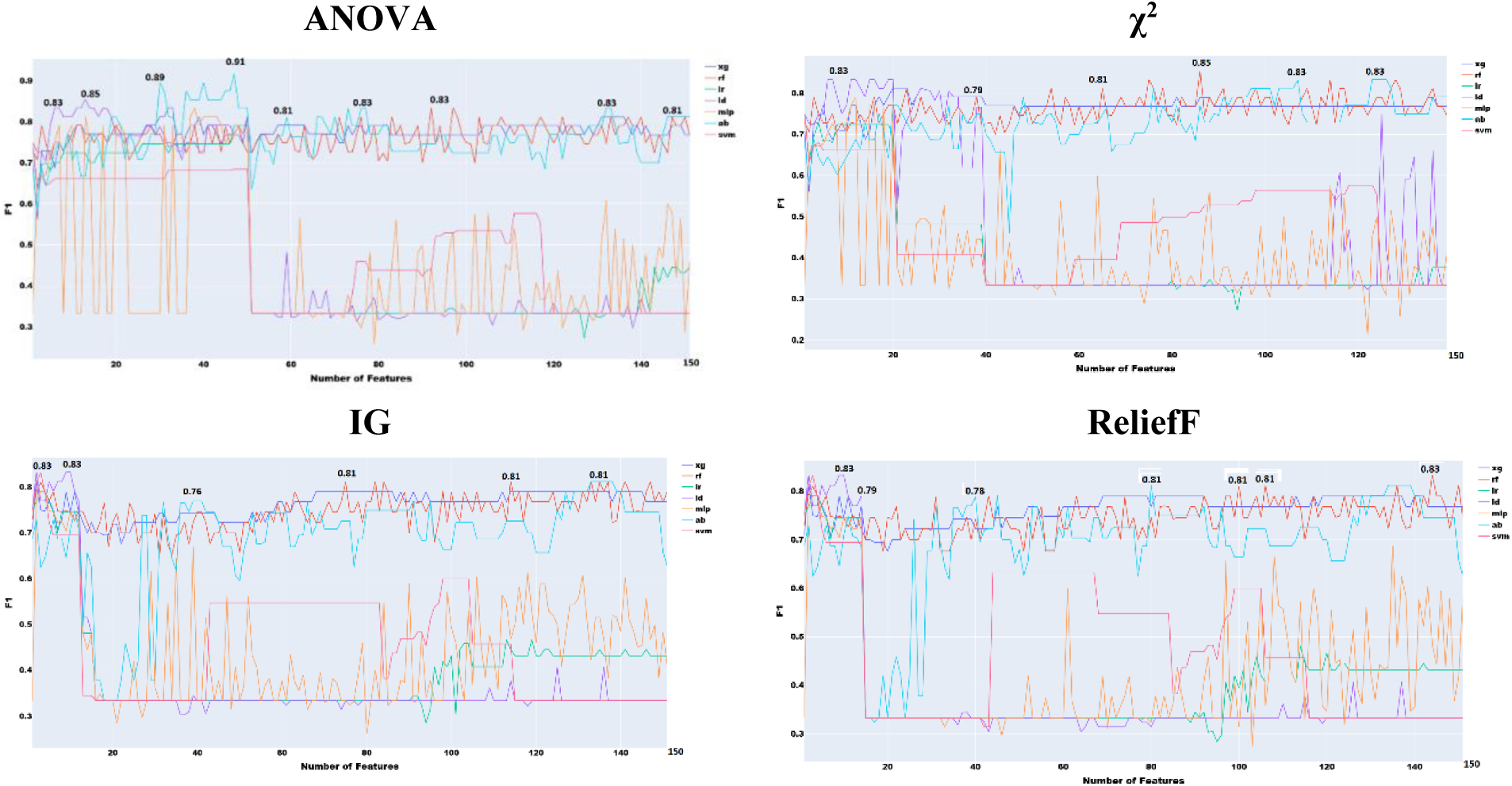
Feature Selection Experiments with Prosodic and Phonological Features

As we see in Figure 8, an overall decrease of performance across the four feature selection approaches, with χ^2^ giving the best performance of 0.83. As expected, this performance is lower than the best performance with human transcription. This decrease in performance is due to a combination of two factors. ASR modules are still not perfect, and their performance is not yet on par with human hearing (Dhanjal, 2024; Rista & Kadriu, 2020). The second factor is that the audio files were not collected in ideal condition, making the job of ASR engines harder.

### 5.4 Experiments with Prosodic and Acoustic Features

This section focuses on evaluating the combined use of acoustic and prosodic features. Since both feature types are inherently acoustic, some degree of redundancy is anticipated. Therefore, we do not expect a significant performance gain from their combination. The experiments involving prosodic features follow the same methodology used for other feature types. Initially, the most relevant features from both acoustic and prosodic sets are selected using the mRMR method. Subsequently, all four previously adopted feature selection techniques are applied to the combined feature set.

As anticipated, combining prosodic and acoustic features did not lead to any performance improvement compared to using them separately, confirming the redundancy between these two feature sets (Figure 9). The highest performance achieved was 0.83, using χ², Information Gain, and ReliefF, which remains significantly lower than the best results obtained with individual feature types.

### 5.5 Experiments with Acoustic and Phonological Features

The objective of this experiment is to explore the potential complementarity between acoustic and phonological features. Unlike acoustic and prosodic features, which share a similar nature and exhibit redundancy as we saw in section 5.4, acoustic and phonological features represent distinct aspects of speech. Acoustic features primarily capture low-level signal characteristics, while phonological features reflect higher-level linguistic information. Due to this inherent difference, we anticipate a stronger complementary effect between these two groups, which could lead to improved performance. To conduct this analysis, we utilized the combined feature set described earlier, referred to as the double feature, which resulted in a total of 383 features being used in the experiments.

As illustrated in Figure 10, the combination of acoustic and phonological features does not lead to any noticeable improvement over the use of either feature set independently. In fact, the highest performance achieved with the combined feature set is 0.87, which is identical to the best result obtained using only phonological features with roughly the same number of features (Figure 9). This outcome indicates that, despite the theoretical differences between acoustic and phonological features, their practical complementarity is limited in this context. The lack of performance gain suggests that the information captured by each set may overlap more than initially expected, or that the current feature selection and modeling approach may not be effectively leveraging the differences between the two.

### 5.6 Experiments with Prosodic and Phonological Features

In this experiment, we investigate the effectiveness of combining two subgroups of features: phonological and prosodic. These feature types differ fundamentally in nature, phonological features are typically discrete, representing symbolic or categorical information about the sound structure of speech (such as phonemes or syllables), whereas prosodic features are continuous, capturing dynamic aspects of speech such as pitch, F0, and amplitude. Despite their differences, both feature sets relate to speech rhythm and structure, and may offer complementary information.

Prosodic features, in particular, can reflect prosodic characteristics indirectly. For example, features like speech rate, measured in words per second, syllables per second, or phonemes per second, span across multiple linguistic domains, integrating acoustic, phonological, and even morphological cues.

In total, this combined feature set consists of 153 features, initially obtained using mRMR. The features were then ranked using our four adopted selection methods to determine their relative importance. This ranking helps reveal which features, across both prosodic and phonological domains, contribute most significantly to performance and highlights how cross-domain features such as speech rate and turn length may bridge gaps between linguistic levels.

As seen in Figure 11, the best overall performance (F1=0.87) is achieved using the four feature selection approaches with the combination of prosody and phonology. This performance is the same achieved by prosody and phonology individually. This suggests that combining both types does not lead to improvement.

### 5.7 Experiments Using the best of Phonological, acoustic, and Prosodic features

The combination of all the speech features used in the above sections within one set of features can help identify an optimal combination of features that can lead to the best performance. Hence the experiment.

As illustrated in Figure 12, χ² feature selection yielded the highest F1 score (0.89), outperforming all other conventional machine learning approaches. Using logistic regression, this method benefited from the broad coverage of prosodic, acoustic, and phonological features, enabling robust AD detection from speech. This performance was also obtained using χ^2^ and soft voting ensemble learning but with 227 features only (less than the 256 used by the classifier built with acoustic features only). Notably, this performance is equivalent to that of the ensemble model trained on acoustic features alone.

**Figure 12.**
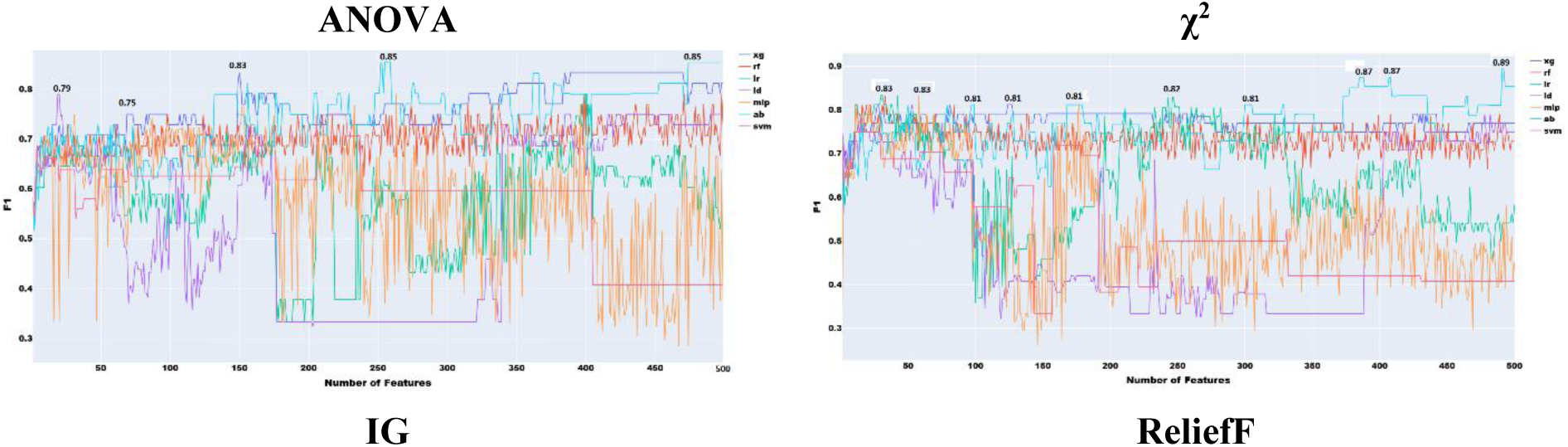
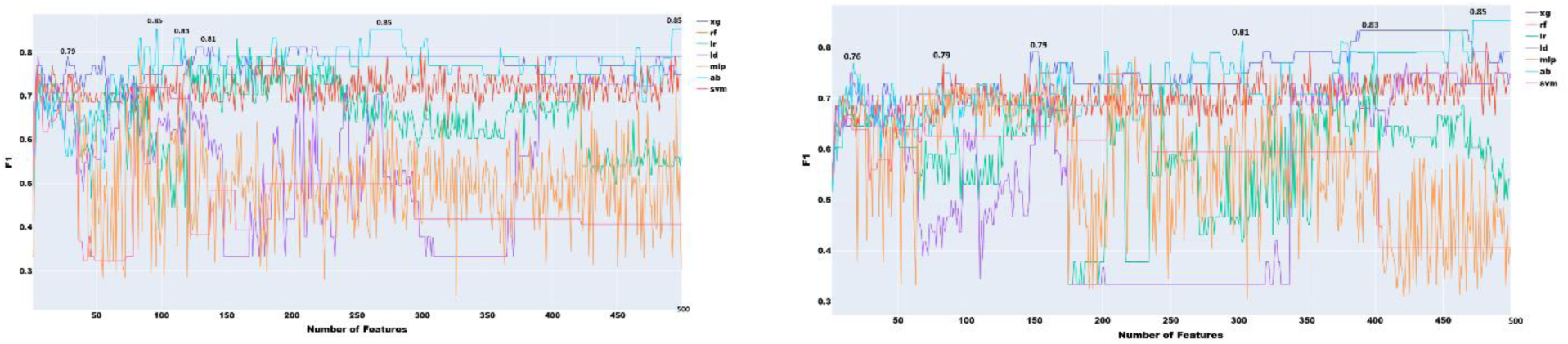
Feature selection experiments with all the speech features combined

The confusion matrix (Table A24) illustrates the top classifier’s performance, which employed soft voting ensemble learning with χ²-selected features. The model accurately identified 20 AD cases and 23 non-AD cases, producing only one false positive but missing four AD cases, indicating a conservative bias that minimizes false positives. While this results in slightly lower recall, there is potential to capture more AD cases without compromising specificity.

Although the F1 score is marginally below the 0.91 reported by Kurdi (2024a) using lexical features, it remains near state-of-the-art for models relying solely on automatically extracted speech features. These findings emphasize the diagnostic value of acoustic, prosodic, and phonological cues and underscore the importance of careful feature selection in early dementia detection.

### 5.8 Feature Ranking

To discuss the contribution of individual features that contributed to the top performance shown in Figure 12, a SHAP diagram is provided in Figure 13.

**Figure 13.**
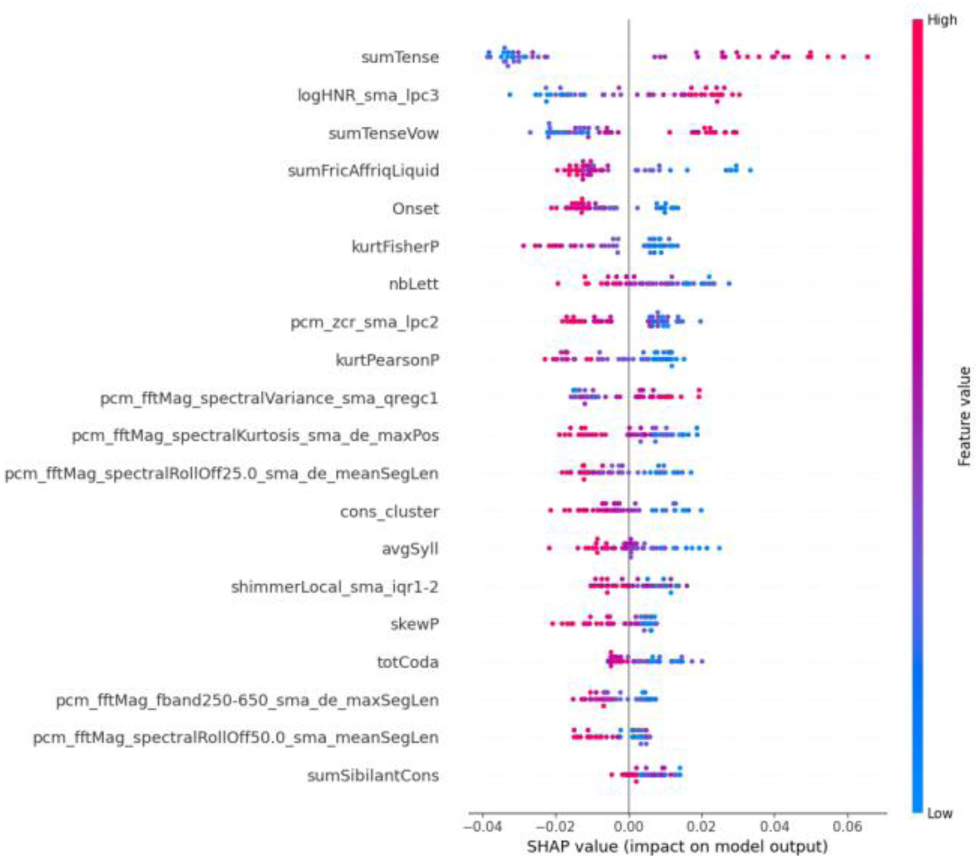
SHAP plot showing the top 20 most important features based on the classifier presented in Figure 12

The SHAP summary plot (Figure 13) reveals that the top twenty features encompass a diverse mix of phonological (e.g., sumTense and cons_cluster), acoustic (e.g., logHNR_sma_lpc3 and pcm_fftMag_spectralVariance_sma_qreg1), and prosodic measures (e.g., shimmerLocal_sma_iqr1-2).

This variety indicates that the model effectively incorporates information from multiple speech domains. The presence of features from different areas, with no single domain dominating the predictions, highlights that the model utilizes complementary cues across speech dimensions. This validates our decision to use a comprehensive feature set, which contributes to the model’s robust overall performance.

The small number of false negatives is notable, but further improvements could be made by expanding training data or incorporating multimodal inputs. Overall, the model shows promise as a non-invasive, speech-based screening tool.

### 5.9 Generalization Experiments Using the University of Delaware’s MCI Dataset

To evaluate the generalizability of our model trained on the ADReSS Challenge dataset, we tested it on the University of Delaware (UD) corpus, which includes speech from individuals with Mild Cognitive Impairment (MCI), a related but less severe neurodegenerative condition. Unlike the ADReSS dataset, which contains only Cookie Theft picture descriptions, the UD corpus spans diverse elicitation tasks, including multiple picture descriptions (Cookie Theft, Cat Rescue, Going and Coming), story narratives (Cinderella), procedural narratives (peanut-butter-and-jelly sandwich), and personal narratives (Hometown). This task heterogeneity poses a substantial test of robustness, reflecting the difficulty of applying a model trained on a single, uniform task to more variable and ecologically complex speech. To maintain comparability with ADReSS, a 70–30 train–test split was used for the UD data.

As shown in Table 1, performance declines when models trained on Cookie Theft AD speech are tested on the more varied UD corpus. This drop underscores the challenge of transferring models across disorders (AD → MCI) and across task types. Because AD typically produces more pronounced impairments than MCI, F1 scores for detecting MCI are consistently lower than for detecting AD, though the results still demonstrate that the model captures milder deficits to a meaningful extent. Phonological and acoustic features generalize best, whereas prosodic features perform poorly, likely due to their sensitivity to task structure and individual speaking style.

**Table 1.** Results of the Generalization tests using the UD corpus.

| Linguistic Area | Type | Best F1 MCI Model | Best F1 AD Model |
| --- | --- | --- | --- |
| Phonology | Simple | 0.69 | 0.61 |
| Acoustics | Simple | 0.67 | 0.594 |
| Prosody | Simple | 0.5 | 0.39 |
| All | Simple | 0.65 | 0.60 |
| All | Ensemble | 0.74 | 0.64 |

Combining all feature types improves performance, and the ensemble model yields the strongest results (F1 = 0.74 for MCI; 0.64 for AD), indicating that integrating complementary linguistic and acoustic information enhances robustness.

Overall, these findings show moderate cross-dataset generalizability while underscoring the limitations of training on a narrowly defined elicitation task when broader clinical applicability is desired. The results further confirm that the model trained on the full feature set provides superior performance relative to more restricted feature subsets like the one trained on acoustic features only that has a similar F1 performance of 0.89.

### 5.10 Generalization Experiments Using the Ronald Regan Dataset

The Reagan dataset was used to evaluate how well our AD model, trained on the ADReSS Challenge data, generalizes to a different AD-related corpus. This dataset introduces several notable challenges. First, people with Alzheimer’s may show fluctuating speech patterns, with periods in which cognitive symptoms are less evident. Second, the recordings were produced under inconsistent conditions, leading to differences in audio quality, environment, and microphone setup. Together, these factors add realistic variability that makes the dataset both difficult to model and valuable for assessing speech-based detection of cognitive decline. Because of the limited size of this dataset, only the results using all the features will be presented (Table 2).

**Table 2.** Results of the Generalization tests using the Regan corpus using all the features.

| Best F1 Regan’s data Model | Best F1 Adress data Model |
| --- | --- |
| 0.77 | 0.61 |

The model obtained an F1 score of 0.77 on the Reagan dataset, exceeding its performance on the ADReSS test set (F1 = 0.61). This result suggests that the model generalizes reasonably well to an external corpus despite substantial variability in recording conditions and speech quality. This performance may be attributable to a pronounced phonological, prosodic and acoustic manifestations of cognitive decline in Reagan’s later speeches, which align with the patterns learned from the ADReSS training data.

## 6. Discussion and Limitations

### 6.1 General Discussion

To contextualize our findings, we compared our results with prior studies using the ADReSS Challenge dataset, allowing a direct evaluation of speech-based AD detection performance.

Luz et al. (2020) used a comprehensive acoustic feature set, including MFCCs, jitter, shimmer, and harmonic-to-noise ratios, achieving an F1 score of 0.62 with speech features alone. Martinc and Pollak (2020) applied clustering to MFCCs and eGeMAPS features with audio duration and linguistic embeddings, reporting 0.77 accuracy. Balagopalan et al. (2020) combined speech and semantic features, reaching F1 = 0.84 with an SVM classifier. Cummins et al. (2020) explored Bag-of-Audio-Words representations and deep learning models, achieving F1 scores between 0.62 and 0.73 depending on features and architecture.

Other notable approaches include Pappagari et al. (2020), who leveraged x-vector embeddings from speaker recognition (up to 0.66 accuracy), Edwards et al. (2020), who integrated OpenSMILE, GeMAPS, and ComParE acoustic descriptors with phonemic data (F1 = 0.78), and Sarawgi (2020), who focused on disfluency and low-level descriptors, achieving 0.83 accuracy.

Our findings indicate that ensemble models leveraging acoustic information, together with the model based on the integration of acoustic, prosodic, and phonological features, outperform previously reported speech-only approaches, highlighting the value of our systematic methodology that explores multiple feature sets and machine-learning algorithms. Evaluation on the Delaware corpus confirmed that combining the three types of speech information enhances generalization, as it achieved the best result.

Recent work has emphasized that the clinical value of speech-based digital biomarkers lies not only in classification performance but also in their robustness and longitudinal stability. Spilka et al. (2024) demonstrated that a speech-derived biomarker from Clinical Dementia Rating Clinical Dementia Rating (CDR) interviews exhibited moderate test–retest reliability (ICC ≈ 0.62) and tracked clinical progression across ADAS-Cog, CDR-SB, and ADCS-ADL, despite not reporting traditional machine learning metrics such as F1 or accuracy. This framework underscores the importance of external validity, longitudinal sensitivity, and clinical correlation in biomarker evaluation. In this context, our cross-cohort generalization results on the Delaware and Reagan datasets provide initial evidence of robustness across different recording conditions and populations. Future studies should build on this work by incorporating test–retest reliability and longitudinal cognitive or biomarker trajectories, in line with established digital biomarker validation standards.

### 6.2 Clinical Discussion

These findings have important clinical implications for the early detection and monitoring of Alzheimer’s disease (AD). Speech is a natural, accessible, and low-burden biomarker that can be collected unobtrusively in primary care or telehealth settings. Our results align with evidence that acoustic, prosodic, and phonological changes emerge early in cognitive decline, often preceding deficits detected by standard neuropsychological assessments.

Models that integrate multiple speech-feature types performed strongly, reflecting AD’s impact across rhythm, articulation, and spectral characteristics. Capturing this multidimensional profile may enable more sensitive and comprehensive screening tools, complementing traditional cognitive tests and supporting earlier identification of at-risk individuals.

The interpretability of our feature-selection approach further enhances clinical relevance by highlighting which speech characteristics most influence model decisions, fostering clinician trust and linking specific speech alterations to underlying neurocognitive processes.

Speech-based assessments are noninvasive, low-cost, and easy to administer, offering opportunities to expand cognitive screening, particularly in underserved or remote populations. Reliance on non-lexical speech features also reduces dependence on language proficiency, education, and cultural context, mitigating biases common in conventional cognitive tests. Combined with passive collection via mobile or telephone platforms, these features support scalable, remote screening within routine clinical workflows and community programs.

Although our findings are favorable relative to prior ADReSS studies, clinical translation requires validation in larger, longitudinal, and more diverse cohorts. Future research should examine longitudinal trajectories of these speech features and their associations with established AD biomarkers, including neuroimaging, cerebrospinal fluid measures, and neuropsychological performance.

### 6.3 Limitations and Future Work

A primary limitation of this study is its reliance on the ADReSS Challenge dataset. Although this dataset is balanced and widely used for benchmarking, it remains relatively small, demographically homogeneous, and collected under controlled conditions using a single narrative task (the Cookie Theft picture description). These constraints may limit ecological validity and reduce the generalizability of our findings to more diverse, multilingual, or real-world populations. Future work should therefore validate the proposed approach on larger and more heterogeneous datasets that include multiple narrative tasks, spontaneous speech, and variable recording environments.

The dataset’s cross-sectional design further restricts the analysis to binary classification of Alzheimer’s disease (AD) versus healthy controls, preventing investigation of disease progression, early detection among at-risk individuals, or longitudinal patterns of speech decline. Future studies should incorporate longitudinal datasets to monitor speech changes over time and relate them to established biomarkers such as neuroimaging, cerebrospinal fluid measures, and standardized cognitive assessments.

This study focuses exclusively on speech, enabling an in-depth examination of acoustic, prosodic, and phonological features in AD. While this provides valuable insight into spoken-language changes, speech represents only one dimension of the broader linguistic impairments associated with AD. This work is part of an ongoing research program: our previous study analyzed lexical features (Kurdi, 2024a), and forthcoming work will address syntax, morphology, and discourse-level phenomena. Integrating these linguistic layers with speech-based models will support a more comprehensive understanding of language impairment in AD.

Although our datasets lack PET/CSF measures, the convergence between our top features and biomarker-associated patterns reported in prior work strengthens the mechanistic plausibility of speech-based detection. Integrating imaging or fluid biomarkers in future work will be essential to determine whether these feature clusters represent valid surrogate indicators of underlying AD pathology.

Another limitation is that the study includes only AD and cognitively healthy controls. In clinical settings, differential diagnosis involves distinguishing AD from other neurological, psychiatric, or cognitive disorders that also affect language (e.g., depression, stress, schizophrenia). Expanding datasets to include such conditions would enhance real-world applicability.

Cognitive fluctuations within individuals with AD may also have contributed to variability in the samples. Moreover, the Reagan dataset used for generalization consists of a single speaker with heterogeneous recordings. Although the model demonstrated robust generalization performance, future work should evaluate multi-speaker datasets and incorporate methods to handle intra-individual variability, including repeated measures and longitudinal designs.

## 7. Conclusion

In conclusion, this study validates a systematic, feature-integrated approach for diagnosing Alzheimer’s disease (AD) from speech, achieving high predictive performance. An F1 score of 0.89 was attained either by ensemble learning on acoustic features alone or by integrating phonological, prosodic, and acoustic features. Integration of multiple feature types yielded the best results on the Delaware corpus used for validation, supporting the methodological advantage of a structured, comprehensive feature set. Feature importance analysis highlighted phonatory instability and temporal disruption as objective acoustic markers of AD. While promising, these findings are based on a controlled dataset; future work should assess their generalizability using longitudinal and ecologically valid speech samples from diverse aging populations.

Looking ahead, the next steps include integrating syntactic, morphological, and discourse-level features with acoustic and prosodic analysis, validating models on larger, multi-task, longitudinal, and multilingual datasets, and expanding classification beyond binary AD versus healthy controls to include other cognitive and psychiatric conditions. Developing robust methods to account for intra-individual variability due to cognitive fluctuations and linking speech features to biological markers and clinical outcomes will further enhance interpretability and translational potential.

Overall, these results reinforce the promise of a multi-feature speech analysis approach as a scalable, low-cost digital biomarker. The methodology is well suited for remote screening within telehealth platforms, supporting earlier, non-invasive detection and timely intervention for individuals at risk of AD, while laying the foundation for combining validated acoustic features with established lexical markers to accelerate clinical adoption.

## 8. Appendix

## Footnotes

1 https://orangedatamining.com/

2 https://pypi.org/project/pymrmr/

3 LPC stands for Linear Predictive Coding, which is a method for representing the digital signal of speech in a compressed form.

## Appendix A. Tables

**Table A3.**
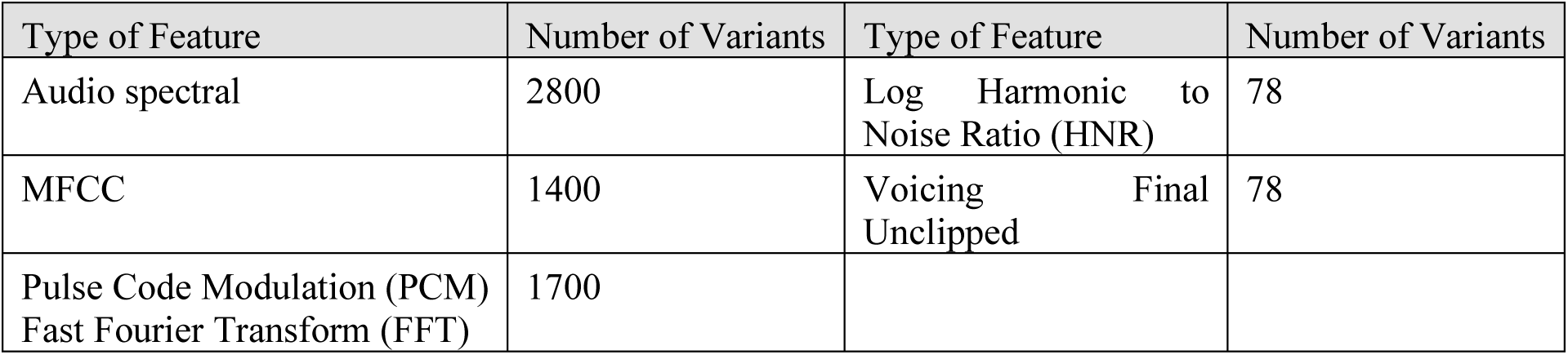
Types and counts of the acoustic features.

| Type of Feature | Number of Variants | Type of Feature | Number of Variants |
| --- | --- | --- | --- |
| Audio spectral | 2800 | Log Harmonic to Noise Ratio (HNR) | 78 |
| MFCC | 1400 | Voicing Final Unclipped | 78 |
| Pulse Code Modulation (PCM)<br>Fast Fourier Transform (FFT) | 1700 |  |  |

**Table A4.** The top 10 audio spectral features.

| Feat | IG | Anova | Chi | Relief |
| --- | --- | --- | --- | --- |
| audSpec Rfilt sma de[21] meanSegLen | A | A | A | A |
| audSpec Rfilt sma[19] maxSegLen | A | A | A | B |
| audSpec Rfilt sma[3] maxPos | A | B | A | B |
| audSpec Rfilt sma de[20] maxSegLen | A | A | A | B |
| audSpec Rfilt sma de[23] meanSegLen | A | A | A | B |
| audSpec Rfilt sma de[17] maxSegLen | A | A | A | C |
| audSpec Rfilt sma de[18] maxSegLen | A | A | A | C |
| audSpec Rfilt sma de[21] segLenStddev | A | A | A | C |
| audSpec Rfilt sma[20] maxSegLen | A | A | B | B |
| audSpec Rfilt sma[2] centroid | A | B | A | C |

**Table A5.** The top 10 MFCC features.

| Feat | IG | Anova | Chi | Relief |
| --- | --- | --- | --- | --- |
| mfcc sma[4] maxSegLen | B | A | B | A |
| mfcc sma[4] segLenStddev | B | B | B | B |
| mfcc sma[14] maxSegLen | A | B | B | B |
| mfcc sma[13] maxSegLen | B | B | B | B |
| mfcc sma[10] maxSegLen | B | B | B | C |
| mfcc sma de[1] meanSegLen | C | B | C | A |
| mfcc sma[13] segLenStddev | C | B | B | C |
| mfcc sma[3] maxSegLen | C | B | C | B |
| mfcc sma[6] maxSegLen | C | B | C | C |
| mfcc sma[7] range | C | D | C | B |

**Table A6.**
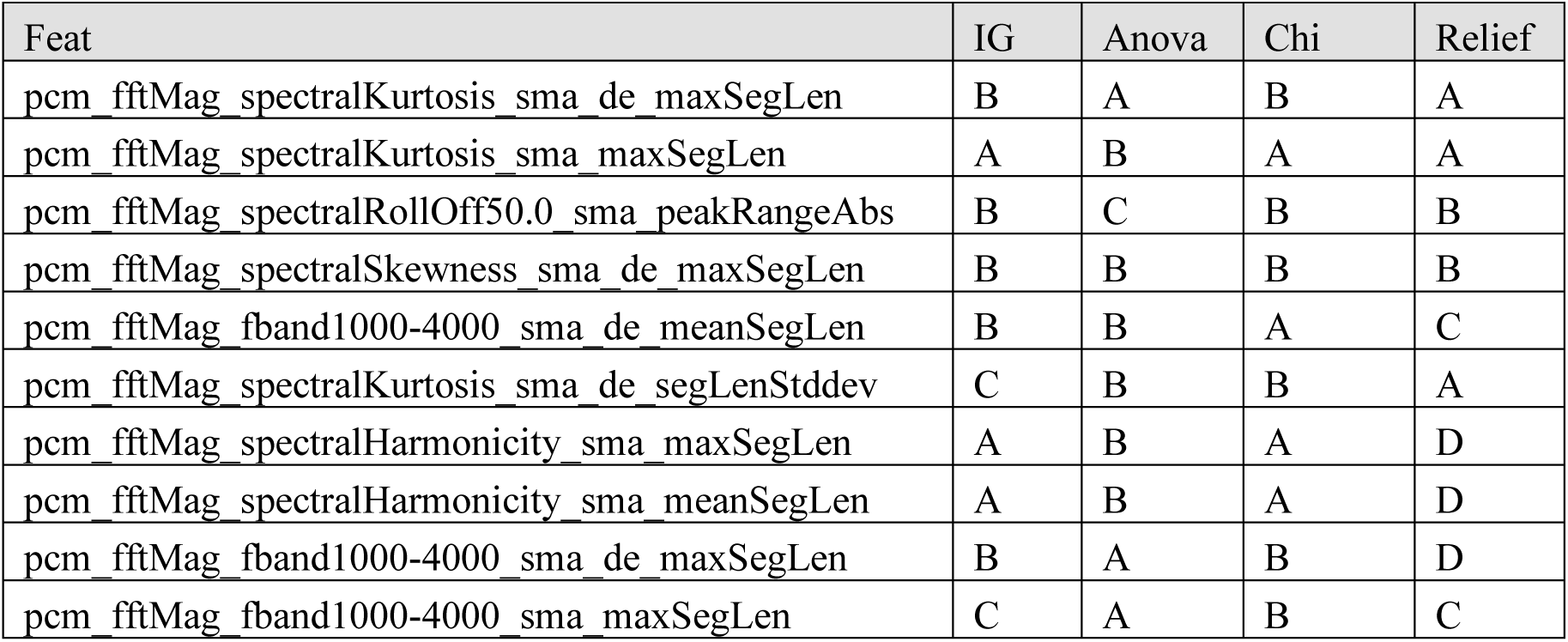
Top 10 PCM features.

**Table A7.** The top 6 LogHNR and voice final unclipped features.

| Feature | IG | Anova | $\chi^2$ | ReliefF |
| --- | --- | --- | --- | --- |
| logHNR sma lpc3 | B | A | B | D |
| voicingFinalUnclipped sma de kurtosis | D | B | D | C |
| voicingFinalUnclipped sma qregc2 | None | None | D | None |
| logHNR sma de kurtosis | D | D | D | None |
| voicingFinalUnclipped sma de leftctime | None | None | None | D |
| logHNR sma de leftctime | D | None | None | None |

**Table A8.**
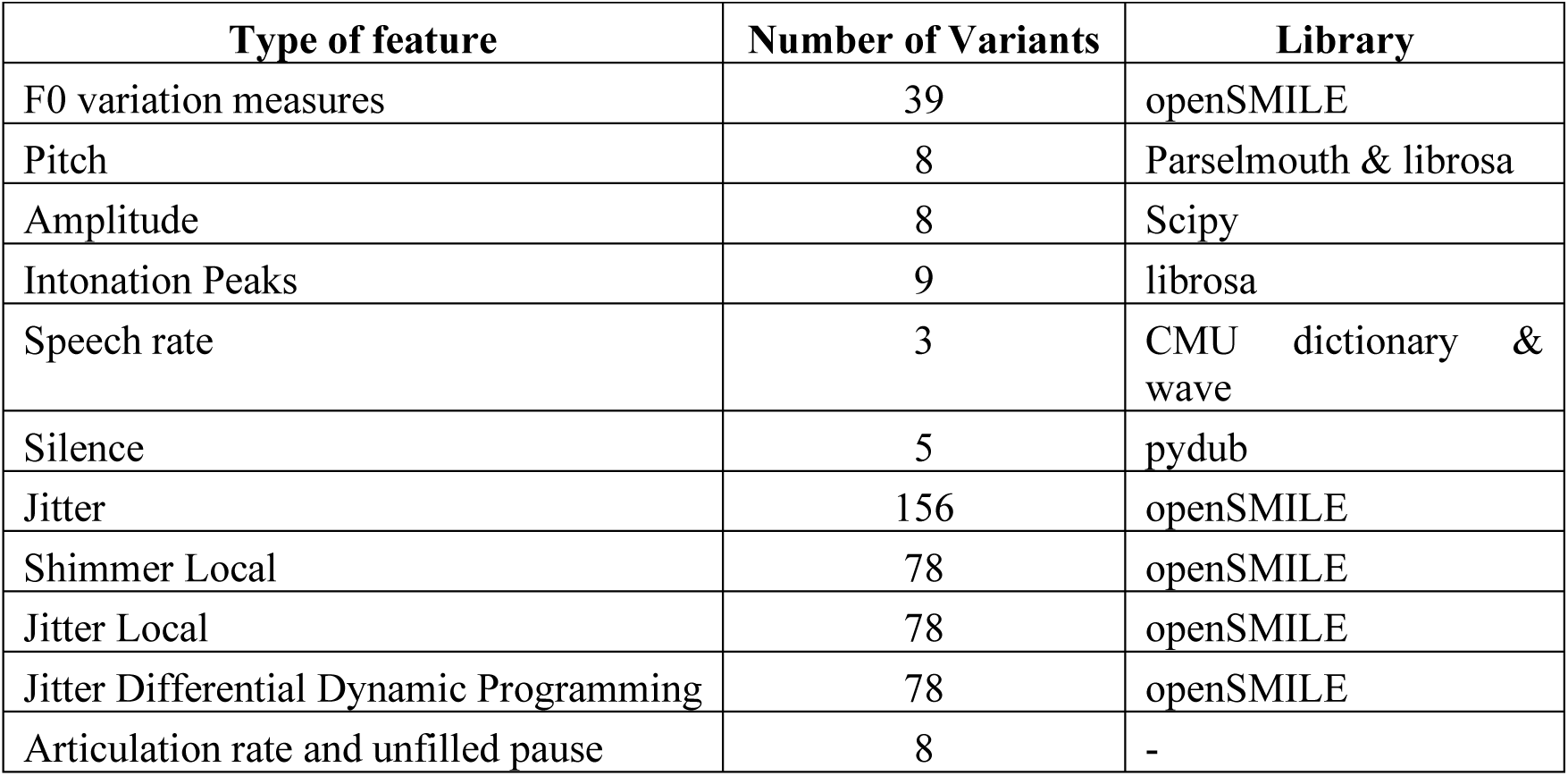
Prosodic features extracted from speech files.

| Type of feature | Number of Variants | Library |
| --- | --- | --- |
| F0 variation measures | 39 | openSMILE |
| Pitch | 8 | Parselmouth & librosa |
| Amplitude | 8 | Scipy |
| Intonation Peaks | 9 | librosa |
| Speech rate | 3 | CMU dictionary & wave |
| Silence | 5 | pydub |
| Jitter | 156 | openSMILE |
| Shimmer Local | 78 | openSMILE |
| Jitter Local | 78 | openSMILE |
| Jitter Differential Dynamic Programming | 78 | openSMILE |
| Articulation rate and unfilled pause | 8 | - |

**Table A9.** Top 10 F0 features.

| Feature | IG | Anova | $\chi^2$ | ReliefF |
| --- | --- | --- | --- | --- |
| F0final sma_lpc2<sup>3</sup> | A | A | A | D |
| F0final sma_maxPos | B | A | A | B |
| F0final sma_lpc1 | A | B | B | C |
| F0final sma_lpc3 | B | A | A | None |
| F0final sma_leftctime | B | A | B | None |
| F0final sma_iqr2-3 | B | B | B | C |
| F0final sma_qregc3 | B | D | A | D |
| F0final sma_minPos | B | D | D | None |
| F0final sma_quartile3 | C | B | B | D |
| F0final sma_risetime | B | None | D | D |

**Table A10.** Top 10 pitch features.

| Feature | IG | Anova | $\chi^2$ | ReliefF |
| --- | --- | --- | --- | --- |
| meanP | A | A | A | A |
| pearModeFirstP | A | A | A | A |
| iqrP | A | A | A | B |
| skewP | A | A | A | D |
| kurtFisherP | A | B | A | D |
| kurtPearsonP | A | B | A | D |
| stdP | B | D | B | A |
| varP | B | D | B | B |
| meanPitchLibroP | B | None | D | D |
| maxPitchLibroP | C | D | D | D |

**Table A11.** Top 10 amplitude features.

| Feature | IG | Anova | $\chi^2$ | ReliefF |
| --- | --- | --- | --- | --- |
| IQR | B | A | B | B |
| Kurt Pearson | D | C | B | B |
| mean | A | D | D | D |
| Pearson Mode First | A | D | D | D |
| Kurt Fisher | D | C | C | C |
| max | D | D | None | A |
| std | None | D | None | A |
| range | D | None | None | A |
| min | None | None | None | A |
| var | None | D | None | B |
| mode | D | D | None | D |
| delta | D | None | D | D |
| Pearson Mode Second | D | D | None | D |

**Table A12.**
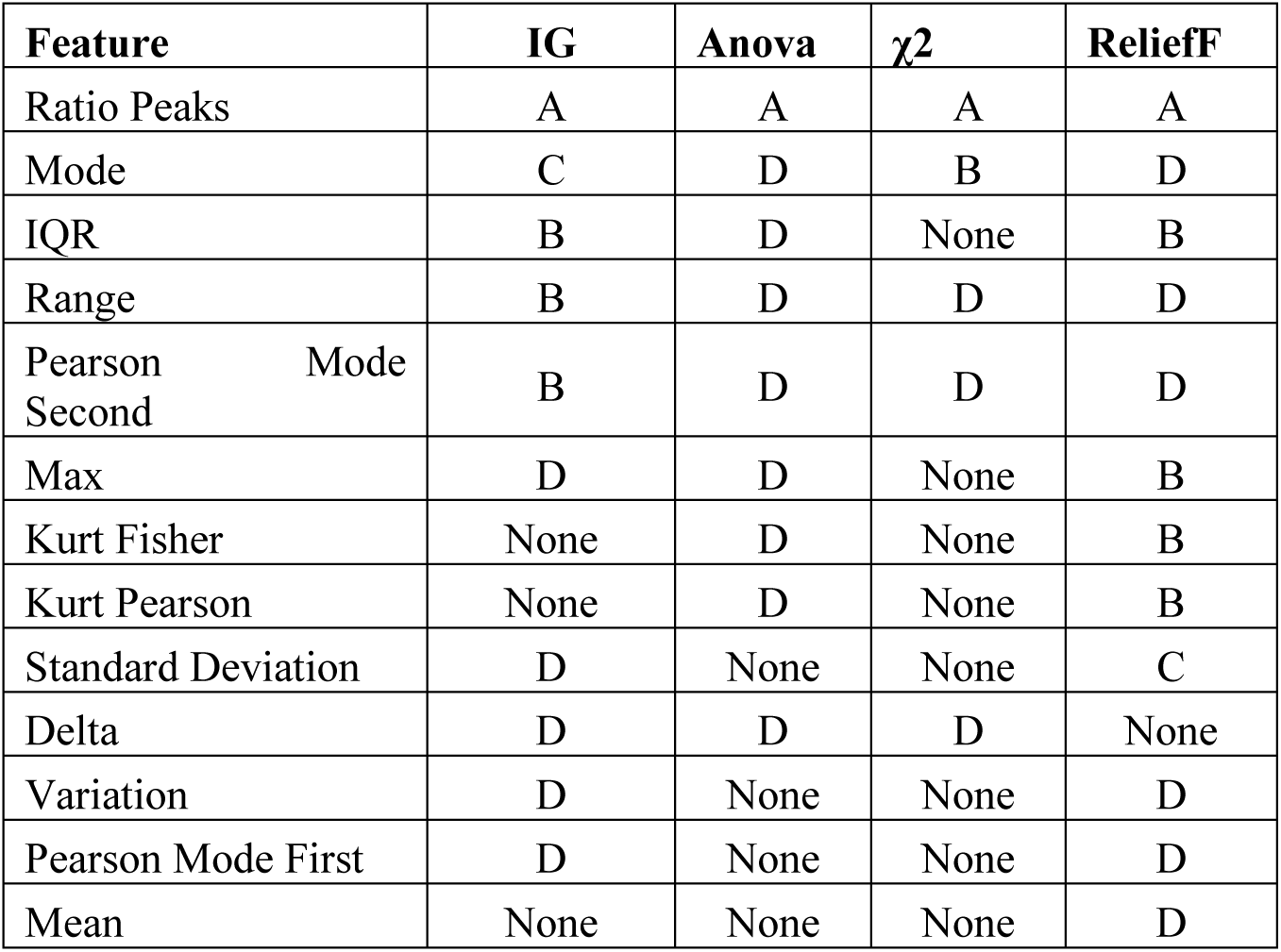
Ranking of the intonation peak features.

**Table A13.** Ranking of the three articulation rate features.

| Feature | IG | Anova | $\chi^2$ | ReliefF |
| --- | --- | --- | --- | --- |
| ratioWords | D | D | D | None |
| ratioChars | D | D | D | None |
| ratioSylls | None | D | D | None |

**Table A14.** Ranking of the unfilled features.

| Feature | IG | Anova | $\chi^2$ | ReliefF |
| --- | --- | --- | --- | --- |
| Ms10 | C | D | B | B |
| Ms200 | B | D | C | None |
| Ms25 | D | D | D | D |
| Ms50 | D | D | D | D |
| Ms100 | D | None | None | D |

**Table A15.**
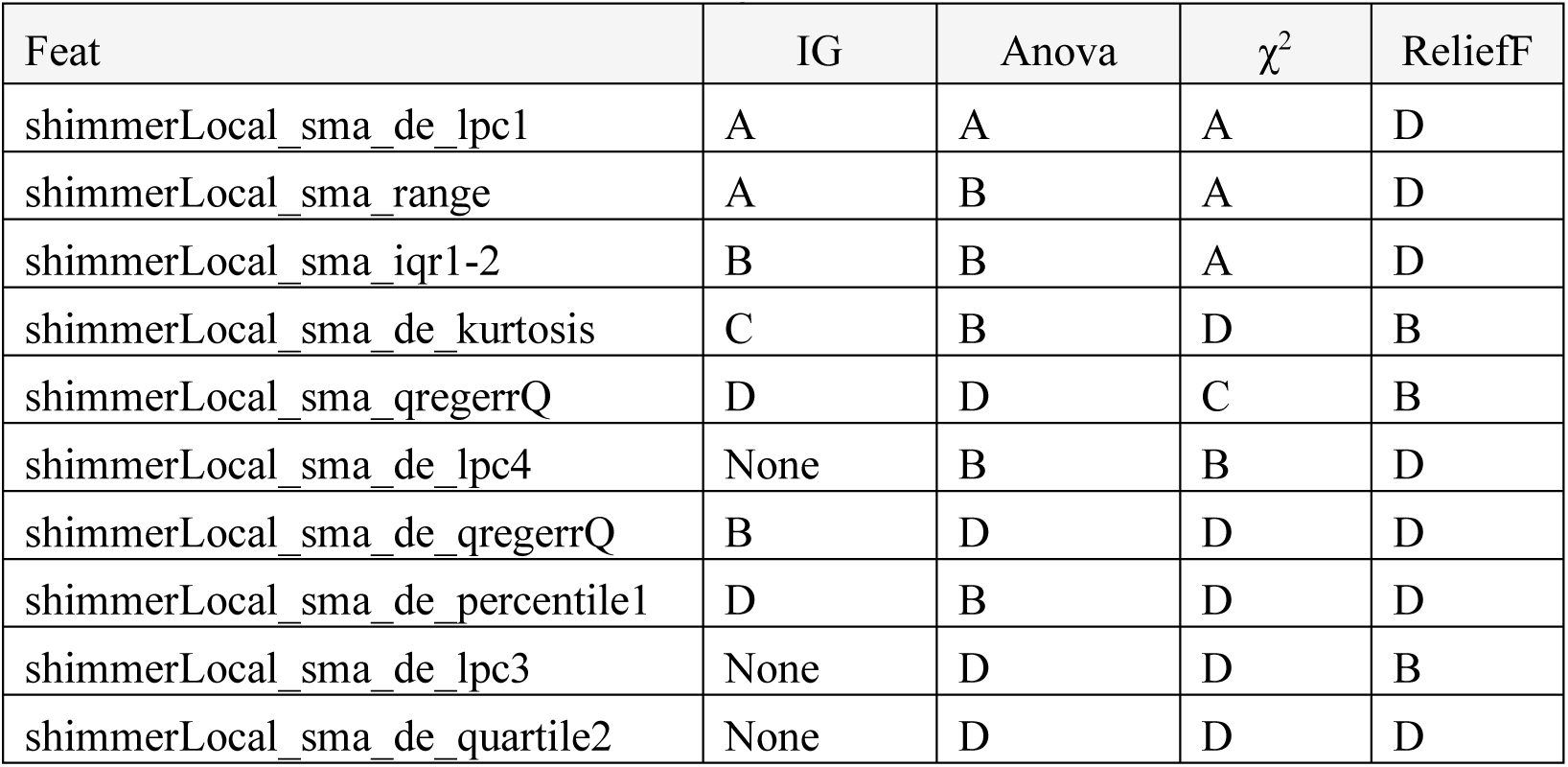
Top 10 shimmer features.

**Table A16.**
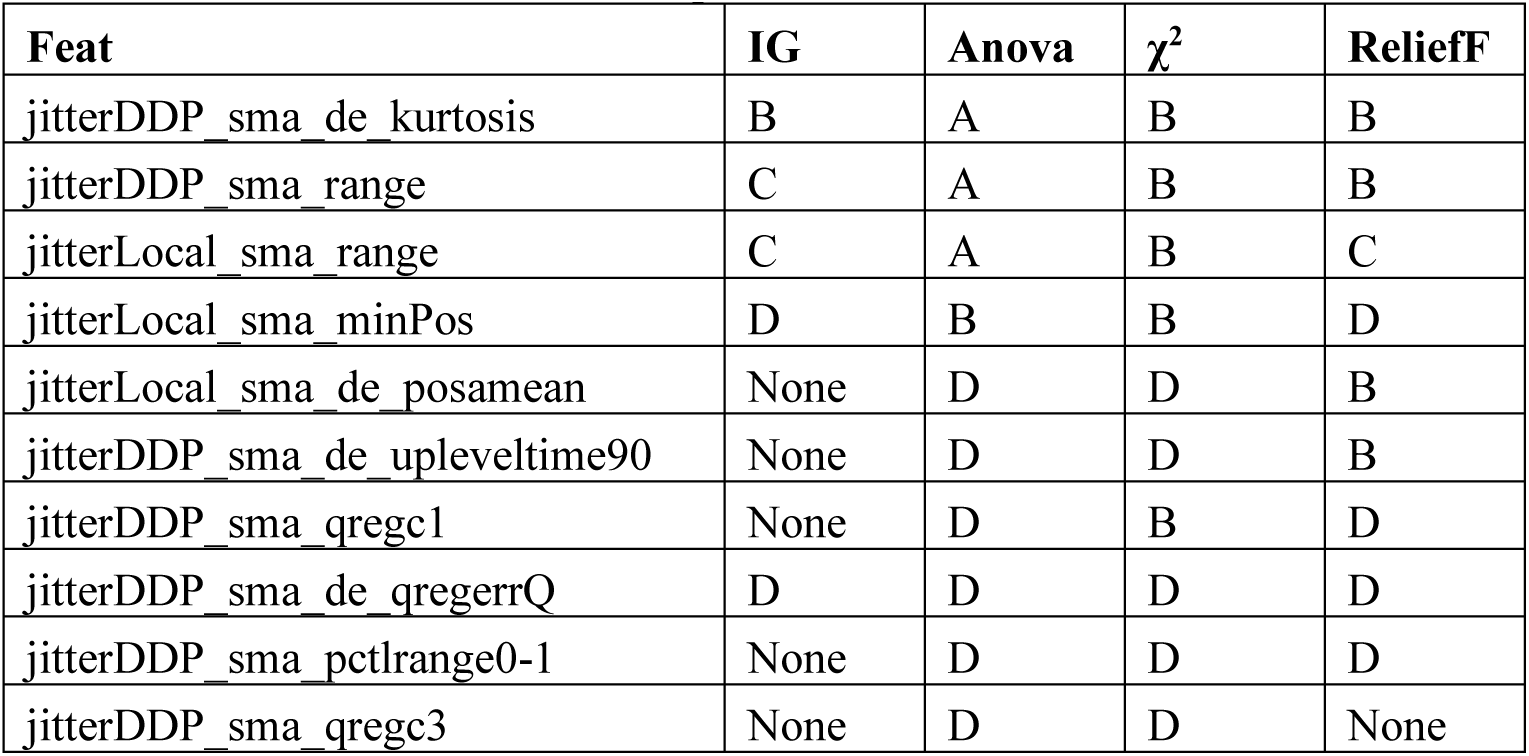
Top 10 jitter features.

**Table A17.** Type of phonetic and phonological features.

| Type | Nb Features |
| --- | --- |
| Consonant Place of Articulation | 22 |
| Consonant Manner of Articulation | 27 |
| Vowel features | 11 |
| Supra-phonemic Structure | 11 |
| Misc. features | 6 |

**Table A18.**
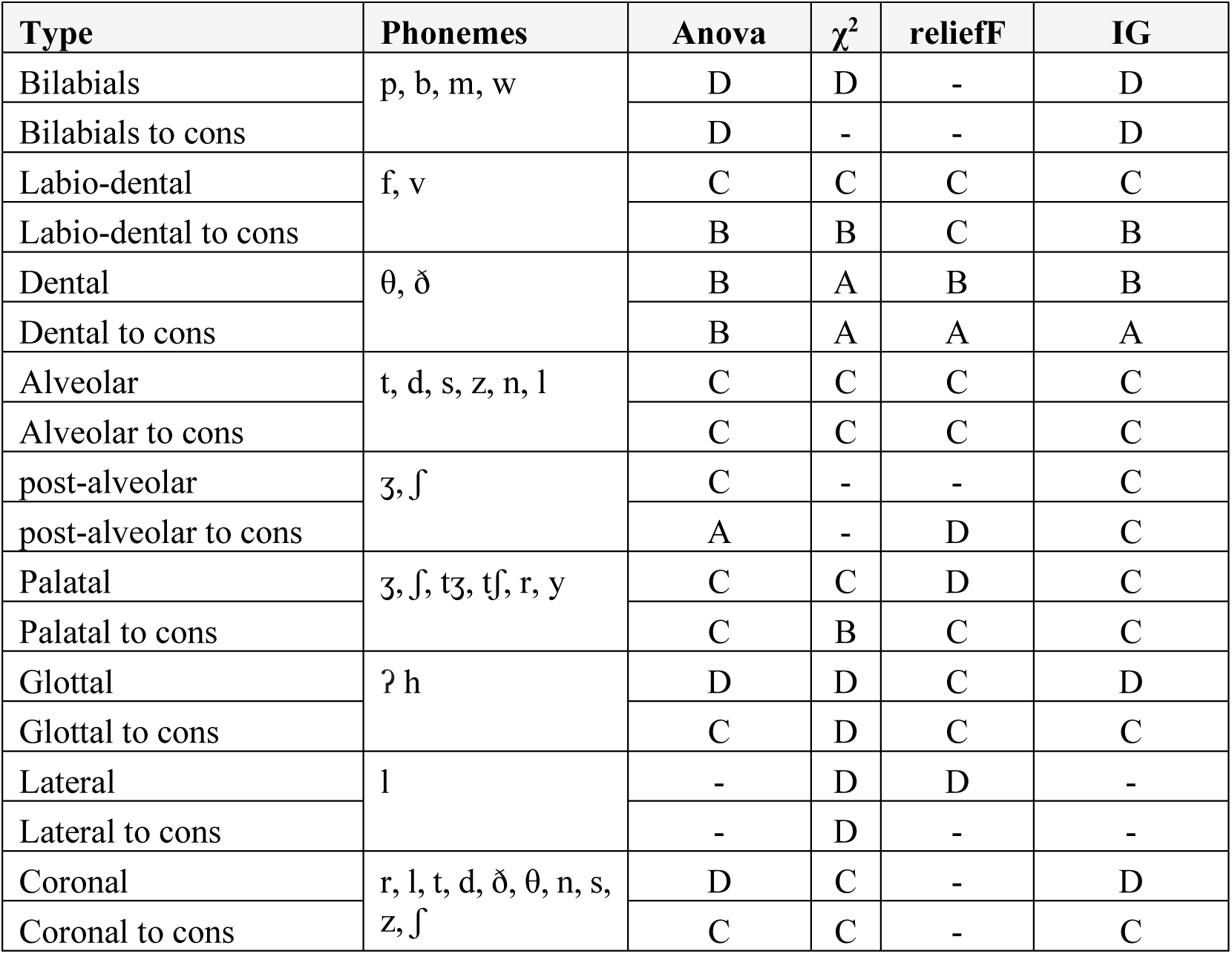
CPA features with their rankings^4^ according to four feature ranking methods: Anova, χ^2^, reliefF, and Information Gain.

**Table A19.** Ranking of CMA features according to the four adopted feature selection methods.

| Type | Phonemes | Anova | $\chi^2$ | reliefF | IG |
| --- | --- | --- | --- | --- | --- |
| Stop | p, b, t, d, k, g, ʔ | - | - | - | - |
| Stop to cons. |  | - | - | - | - |
| Fricatives | v, f, θ, ð, z, s, ʒ, ʃ, h, tʒ, tʃ | B | B | A | B |
| Fricatives to cons |  | A | A | A | A |
| FriAffLiq | v, f, θ, ð, z, s, ʒ, ʃ, t, d, l, r, h | A | B | C | A |
| FriAffLiq to cons |  | A | A | A | A |
| Nasal | m, n, ŋ | D | C | - | D |
| Nasal to cons. |  | - | D | D | - |
| Liquids | l, r | D | C | - | D |
| Liquids to cons. |  | D | C | - | D |
| Glide | w, y | C | B | A | B |
| Glide to cons. |  | B | C | A | B |
| Approximant | w, r, j | C | - | D | C |
| Approximant to cons. |  | - | D | D | - |
| Sonorant | y, w, l, r, m, n, ŋ | - | D | D | - |
| Sonorant to cons. |  | - | - | - | - |
| Continuant | r, l, θ, ð, n, s, ʃ, k, h | D | - | B | D |
| Continuant to cons |  | - | - | C | - |
| Voiced | b, d, g, v, ð, z, ʒ, dʒ, m, n, ŋ, l, r, ɹ, w, j | D | D | D | D |
| Voiced to cons. |  | - | D | D | - |
| Strident | f, v, s, z, ʃ, ʒ, tʃ, dʒ | B | C | D | B |
| Sibilant | s, z, ʃ, ʒ, tʃ, dʒ | C | C | D | C |
| Sibilant to cons. |  | B | B | D | B |
| Distributed | s, z | C | C | D | C |
| Distributed to cons |  | C | C | C | C |
| High | w, ʃ, k, g | - | - | B | - |
| High to cons |  | - | - | C | - |

**Table A20.**
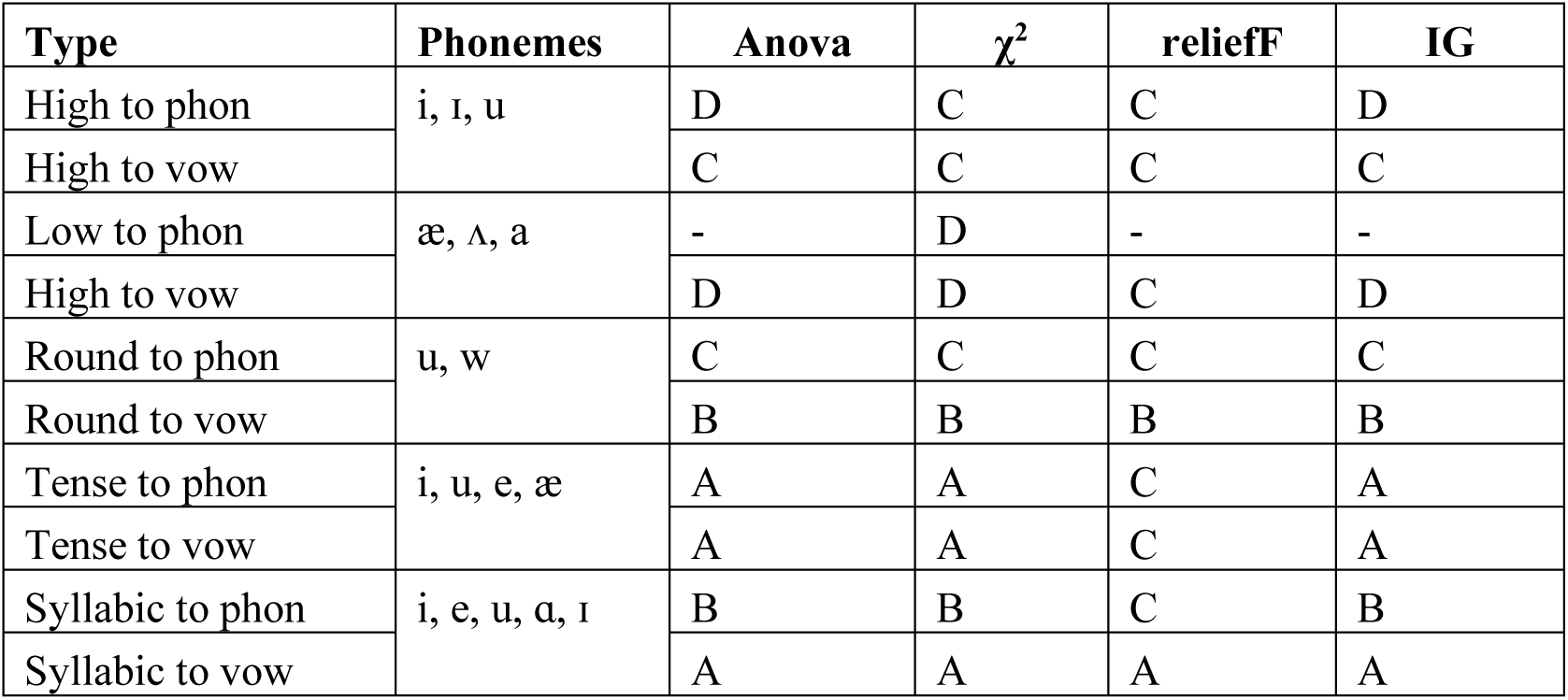
Rankings of the vowels features according to the four adopted feature selection techniques.

**Table A21.** Rankings of Misc. phonemic features according to the four adopted feature selection techniques.

| Feature | Sounds | Anova | $\chi^2$ | reliefF | IG |
| --- | --- | --- | --- | --- | --- |
| Syllabic cons. to phon. | m, n, l | B | B | C | B |
| Syllabic cons. to cons. |  | B | B | B | B |
| Consonantal | r, l, p, t, d, ð, θ, n, s, z, ʃ, k, h | D | D | D | D |
| Diphtongues to phon. | oʊ, aʊ, aɪ, eɪ, ɔɪ, ɪər, ɛər, ʊər | - | - | B | - |
| Diphtongues to cons. |  | D | - | - | D |

**Table A22.**
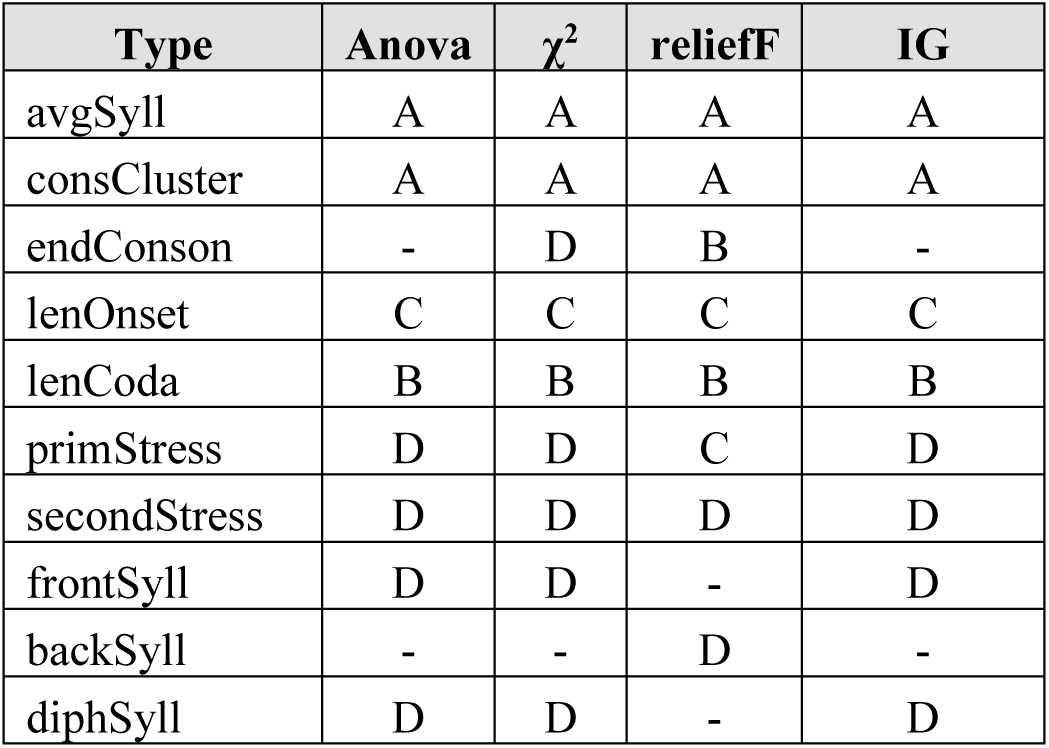
Rankings of the supra phonemic features according to the four adopted features selection methods.

**Table A23.** Confusion matrix of the top classifier with acoustic features.

| Top classifier | F1 | TN | FP |
| --- | --- | --- | --- |
| Feature Selection: Information gain | 0.85 | 20 | 4 |
| Number of features: 256 |  | FN | TP |
| **ML algorithm:** adaptive boosting |  | 3 | 21 |

**Table A24.** Confusion matrices 0.85 two of the top classifiers with prosodic features.

| Top classifiers | F1 | TN | FP |
| --- | --- | --- | --- |
| Feature Selection: Anova | 0.85 | 19 | 5 |
| Number of features: 88 |  | FN | TP |
| ML algorithm: XGBoost |  | 3 | 21 |
| Feature Selection: Anova | 0.85 | TN | FP |
| Number of features: 4 |  | 22 | 2 |
| ML algorithm: AD |  | FN | TP |
|  |  | 5 | 19 |
| Feature Selection: $\chi^2$ | 0.87 | TN | FP |
| Number of features: 51 |  | 19 | 5 |
| ML algorithm: XGBoost |  | FN | TP |
|  |  | 3 | 21 |

**Table A25.** Confusion matrix of one of the top classifiers with phonological features.

| Top classifiers | F1 | TN | FP |
| --- | --- | --- | --- |
| Feature Selection: Anova | 0.87 | 21 | 3 |
| Number of features: 23 |  | FN | TP |
| ML algorithm: XGBoost |  | 3 | 21 |

**Table A26.** Confusion matrix of the top classifier with all the speech features and ensemble learning.

| Top classifier | F1 | TN | FP |
| --- | --- | --- | --- |
| Feature Selection: $\chi^2$ | 0.89 | 23 | 1 |
| Number of features: 227 |  | FN | TP |
| **ML algorithm:** soft voting |  | 4 | 20 |

## Appendix B. Figures

**Figure B14.**
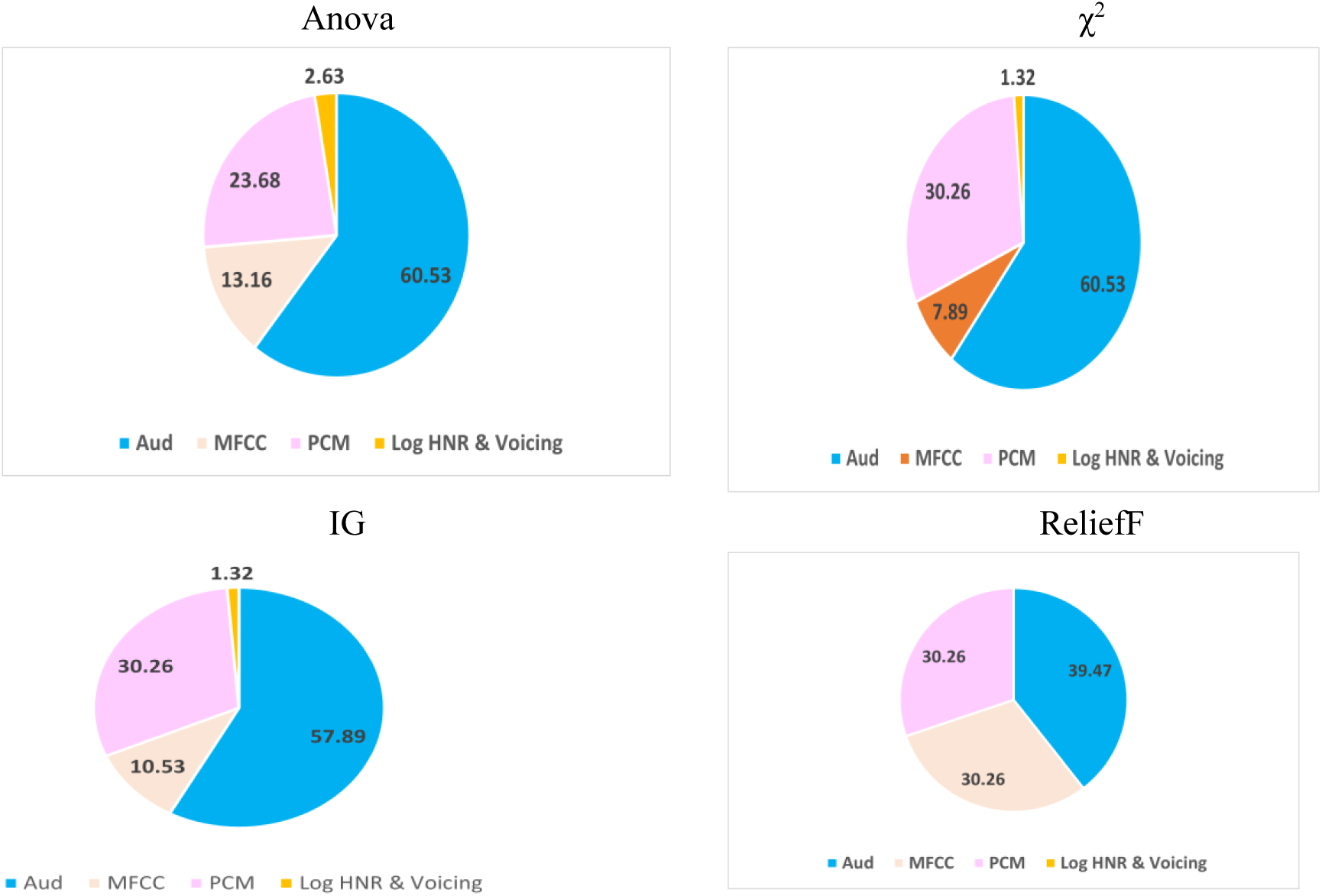
Top 25% of the features according to the five feature ranking methods

**Figure B15.**
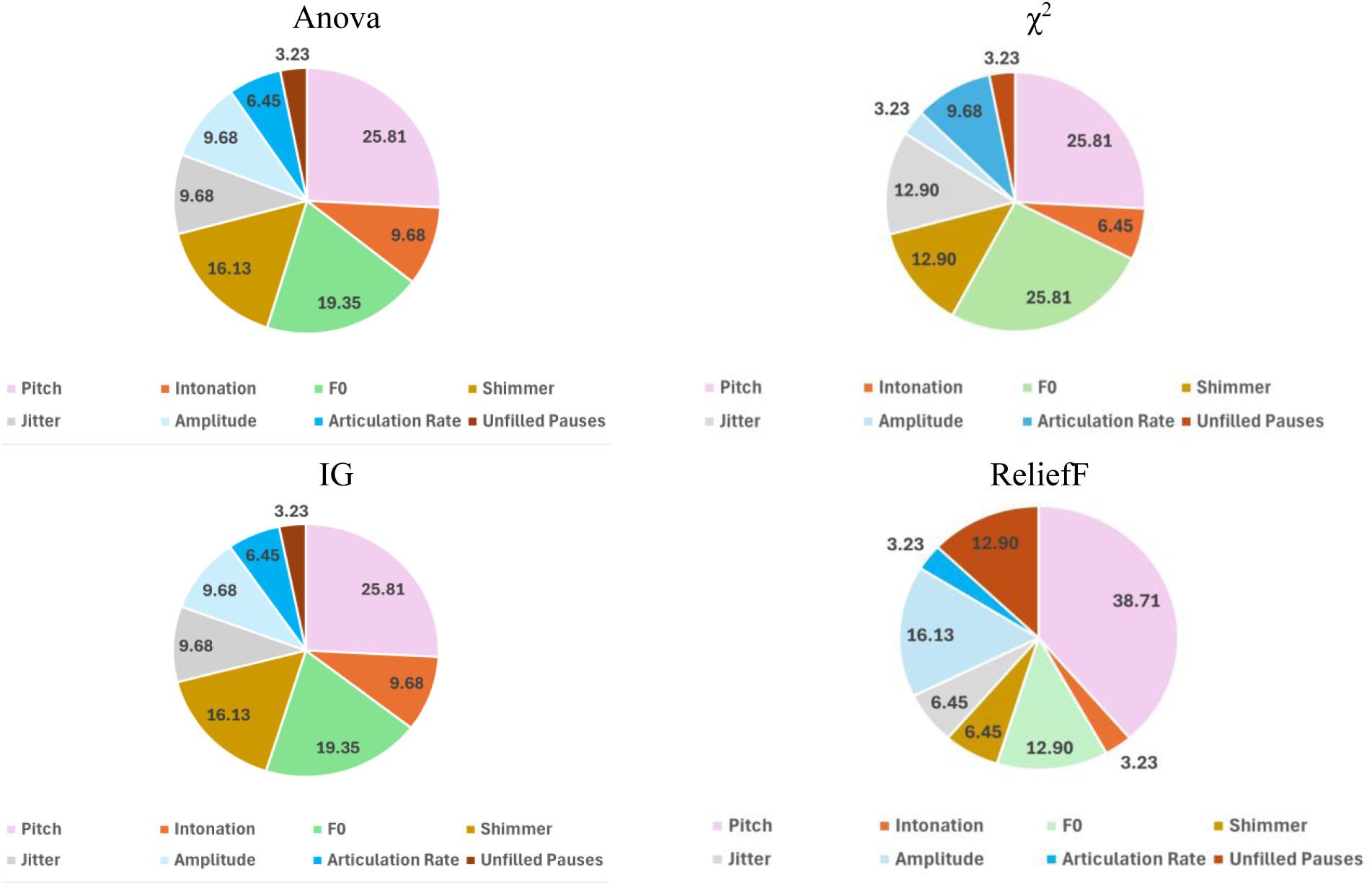
Top 25% of the features according to the five Feature Ranking Methods

**Figure B16.**
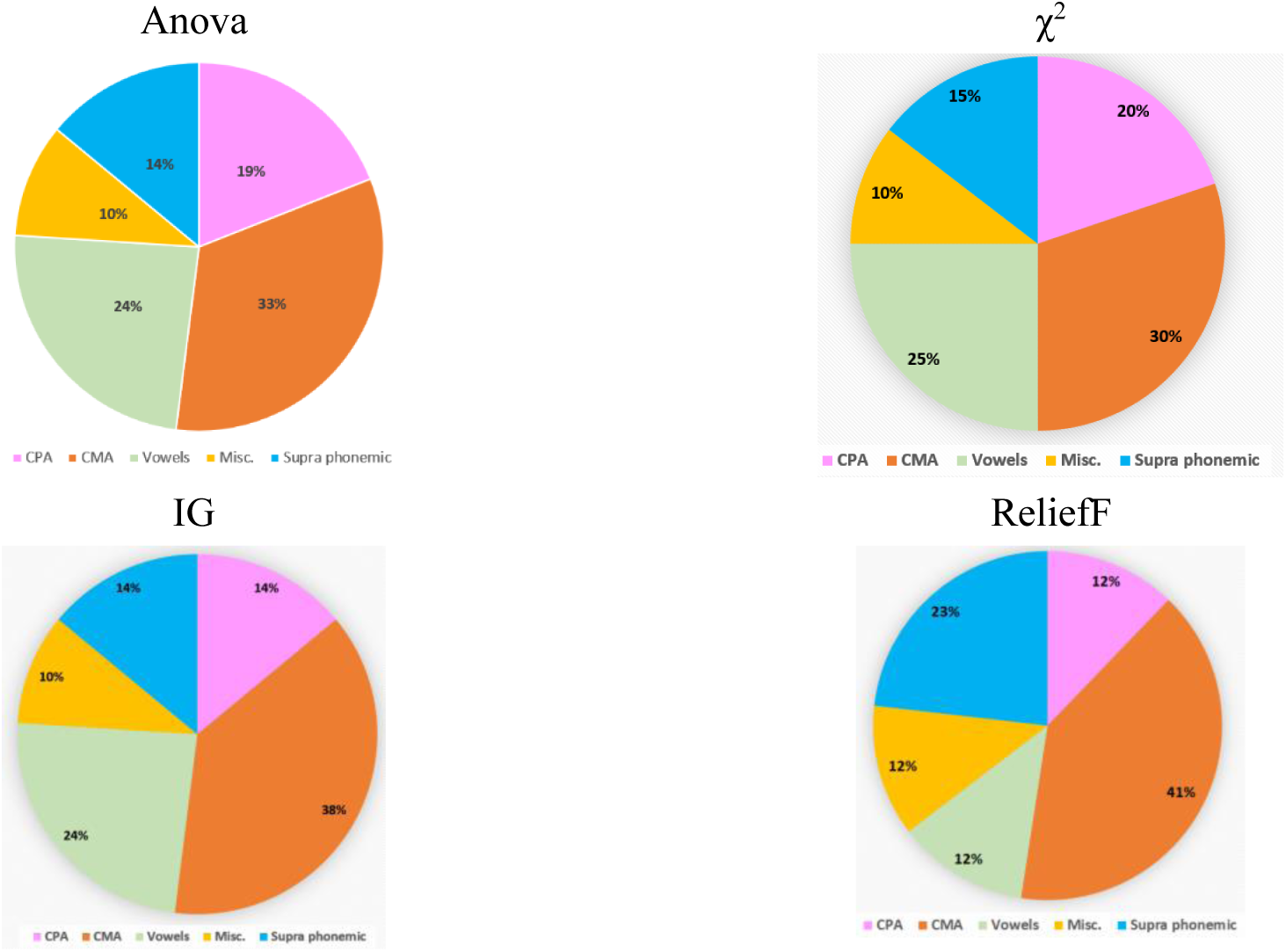
Partition of the top 25% of the phonological features according to the four adopted feature ranking methods

